# Cellular distribution and motion of essential magnetosome proteins expressed in mammalian cells

**DOI:** 10.1101/2023.12.19.572414

**Authors:** Qin Sun, Cécile Fradin, R. Terry Thompson, Frank S. Prato, Donna E. Goldhawk

## Abstract

Magnetosomes are organelle-like structures within magnetotactic bacteria that store iron biominerals in membrane-bound vesicles. In the bacteria, formation of these structures is highly regulated by approximately 30 genes which are conserved throughout different species. To compartmentalize iron in mammalian cells for magnetic resonance imaging using gene-based contrast, we are introducing key magnetosome proteins. We have previously expressed essential magnetosome genes *mamI* and *mamL* as fluorescent fusion proteins in the human melanoma cell line MDA-MB-435 and confirmed their co-localization and interaction. Here we investigate the expression of magnetosome genes *mamB* and *mamE* in MDA-MB-435 cells, using confocal fluorescence microscopy to observe expression patterns and to analyze particle mobility. Custom software was developed to characterize fluorescent particle trajectories. In mammalian cells, essential magnetosome proteins displayed different diffusive behaviours. However, all magnetosome proteins travelled at similar velocities when undergoing directed motion, suggesting that MamL, MamL+MamI, MamB, and MamE interact with similar mammalian mobile elements. These results confirm that localization and interaction of essential magnetosome proteins is tenable in the mammalian intracellular compartment.

## Introduction

In magnetotactic bacteria (MTB), magnetosome formation allows the cell to compartmentalize and concentrate iron biominerals in membrane-enclosed vesicles (1). Magnetosome formation is a stepwise, protein-directed process that begins with vesicle formation and culminates with iron biomineralization. This entire process is regulated by approximately 30 genes, the majority of which are located on the magnetosome genomic island (2, 3). Many of these genes encode membrane proteins which are involved in different steps of bacterial magnetosome formation: inner membrane invagination leading to vesicle formation, recruitment of proteins to the vesicle membrane, alignment of vesicles along a protein filament, concentration of iron inside the vesicle and finally magnetic crystal nucleation and growth. To stimulate the formation of a rudimentary magnetosome-like particle in mammalian cells, we are introducing select genes (alone or in combination) deemed essential for the initial stages of magnetosome formation, namely *mamI*, *mamL*, *mamB* and *mamE*. These four genes are clustered on the *mamAB* operon and are highly conserved in most species of MTB (4). The gene products of *mamI*, *mamL*, and *mamB* are believed to play an essential role in the first steps of magnetosome vesicle formation (2, 5, 6), including designation of the vesicle and recruitment of other magnetosome-associated proteins like MamE (7, 8). The latter plays a crucial role in the initiation of iron biomineralization and recruitment of additional magnetosome-associated proteins (9). We hypothesize that together MamI, MamL, MamB and MamE support the base structure upon which magnetosome formation relies in any cell. As such, we expect these essential magnetosome proteins (co)localize and interact in the intracellular membranous compartment of mammalian systems.

We have previously shown that the red fluorescent MamL fusion protein, Tomato-MamL, assembles into punctate structures that move intracellularly in a mammalian cell line. We have also shown that MamL recruits the green fluorescent MamI fusion protein, EGFP-MamI, to these structures via magnetosome protein-protein interactions. In contrast, EGFP-MamI expressed alone in mammalian cells has a very different cellular distribution (10). The novel and unexpected discovery of the mobility associated with these magnetosome proteins in a foreign cell environment prompted further investigation into particle trajectory analysis.

Here, we quantitatively compare the distribution and mobility of a larger set of magnetosome proteins (MamL, MamI, MamB, MamE), again expressed as fluorescent fusion proteins in a mammalian cell line. When expressed on their own, Tomato-MamL, Tomato-MamB and EGFP-MamE all form (or localize to) punctate structures, which are mostly mobile in the case of MamL and MamB, but immobile in the case of MamE. For EGFP-MamI, co-expression with MamL was necessary to ensure a similar localization to punctate structures so that a comparison in their mobility could be done. These punctate structures will herein be referred to as “particles”. We track fluorescent particle motion and describe their mean-squared displacement (MSD) with a simple power-law analysis to distinguish between different types of mobility. With these tools, the trajectories of punctate intracellular structures are classified into three different categories corresponding to confined, diffusing and actively transported particles. The motions detected for MamL and MamB (and for MamI when interacting with MamL) show that a portion of these structures displays directed motion. This demonstrates the capacity of magnetosome proteins to spontaneously interact with elements of eukaryotic transport machinery.

## Materials and Methods

### Molecular Cloning

Magnetosome genes *mamI* and *mamL* were amplified by PCR from the genomic DNA of *Magnetospirillum magneticum* strain AMB-1 (ATCC 700264) using custom primers (10). Briefly, *mamI* and *mamL* amplicons were purified using a PCR clean-up kit (Invitrogen, Life Technologies, Burlington, Canada); digested with appropriate restriction enzymes; and purified once more, prior to insertion in the cloning vectors pEGFP-C1 (Clontech) and ptdTomato-C1 (Clontech), respectively. All vector-insert constructs used for mammalian cell transfection were propagated in *Escherichia coli* XL10GOLD and verified by sequencing.

Sequence analysis of the MamL protein indicates the presence of a positively-charged peptide (2) in the C-terminal 15 amino acids. Cationic peptides also known as cell-penetrating peptides have been implicated in endocytotic pathways (11). For cloning Tomato-MamL_trunc_, the last 15 amino acids from the C-terminus of MamL were removed by site-directed mutagenesis (10). Briefly, primers were designed to include a stop codon 45 nucleotides upstream from the termination of *mamL*. These primers were used in PCR amplification of the *mamL_trunc_* gene, which was then inserted into the ptdTomato-C1 vector with restriction enzymes EcoRI and BglII.

For cloning FLAG-MamL, primers were designed to flank the *mamL* gene and include an N-terminal FLAG sequence for immunodetection (10). The insert was amplified using PCR, purified using a PCR clean-up kit (Invitrogen, Life Technologies, Burlington, Canada), and restriction digested using SacI and EcoRI. *FLAG-mamL* was then inserted into the pSF-EMCV-*FLuc* vector and propagated in *Escherichia coli* strain XL10GOLD.

For cloning Tomato-MamB, the *mamB* gene was amplified by PCR from AMB-1 genomic DNA using custom primers (Table 1). The amplicon was purified with a PCR clean-up kit, digested with restriction enzymes (Table 1), then purified and inserted into ptdTomato-C1.

**Table 1.**
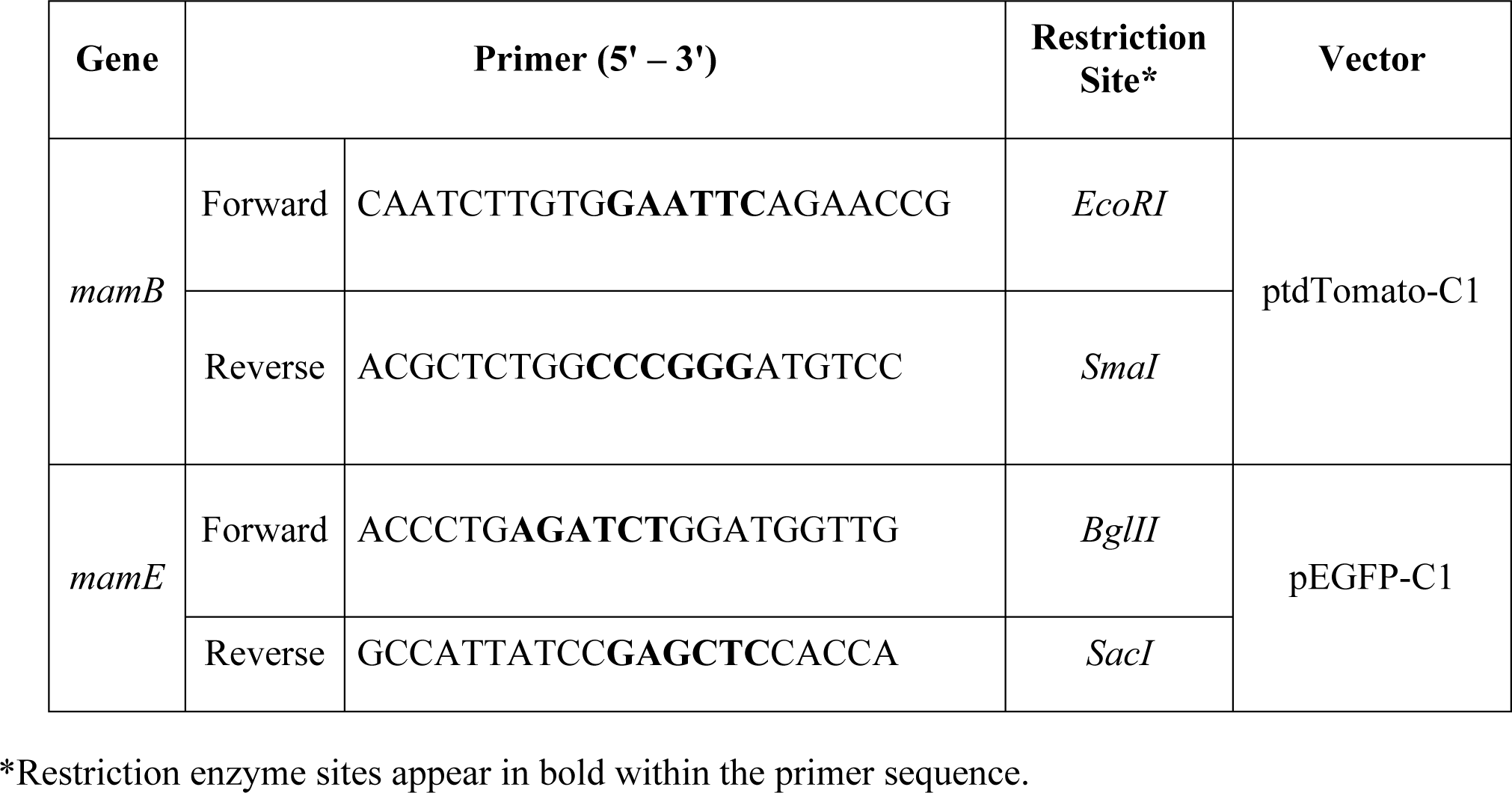
Primer design for cloning MTB genes *mamB* and *mamE*.

For cloning EGFP-MamE, the *mamE* gene was amplified by PCR from the AMB-1 genomic DNA using custom primers (Table 1). The amplicon was purified with a PCR clean-up kit, digested with restriction enzymes (Table 1), then purified and inserted into the pEGFP-C1.

Several attempts at cloning and sequencing EGFP-*mamE* revealed multiple constructs with sporadic mutations relative to the consensus sequence, which itself consists of variable and conserved regions (12) (Supplementary Figure 1). Two of these mutations, denoted EGFP-*mamE* (G49S, T641S) and EGFP-*mamE* (T317A), were used for confocal microscopy and provided comparable results.

### Cell Culture

MDA-MB-435 cells (ATCC HTB-129; derived from an adult female and characterized as a melanoma cell line) are a model of aggressive tumorigenesis (13). Cells were cultured in 100 mm cell culture dishes (CELLSTAR, VWR International, Mississauga, Canada) with Dulbecco’s Modified Eagle Medium (DMEM) containing 1 g/L glucose (Gibco, Life Technologies, Burlington, Canada), 10% fetal bovine serum (FBS; Gibco), 4 U/mL penicillin, and 4 µg/mL streptomycin at 37°C with 5% CO_2_.

To create cell lines expressing red fluorescent fusion proteins tdTomato (Tomato)-MamL, Tomato-MamL_trunc,_ or Tomato-MamB, the enhanced green fluorescent protein (EGFP) fusion protein EGFP-MamE, or the FLAG (DYKDDDDK) tagged protein FLAG-MamL, cells were grown to 60-70% confluency on a 100 mm dish and transfected using Lipofectamine 2000 (Invitrogen), according to company protocol, using 8 µg of each construct. After 16 hours, cells were placed in full medium for 48 hours before commencing antibiotic selection. To select stable cell lines, cells were grown in the presence of 500 µg/mL Geneticin (G418; Gibco) for Tomato or EGFP expression systems and 500 ng/mL Puromycin (Gibco) for the FLAG expression system.

For co-expression of both pEGFP-*mamI* and ptdTomato-*mamL,* cells stably expressing Tomato-MamL were transfected with 8 µg of pEGFP-*mamI*. After 16 hours, cells were placed in full medium for 48 hours before commencing antibiotic selection. Cells were then sorted using fluorescence activated cell sorting (FACS; London Regional Flow Cytometry Facility, Robarts Research Institute, London, Ontario, Canada) to obtain a population of cells fluorescing both green and red (i.e. expressing both EGFP-MamI and Tomato-MamL, respectively). For co-expression with the pSF-FLAG-*mamL*-EMCV-FLuc construct, MDA-MB-435 cells expressing EGFP-MamI were grown to 60-70% confluency on a 100 mm dish and transfected using Lipofectamine 2000 (Invitrogen), according to company protocol, using 8 µg of pSF-FLAG-*mamL*-EMCV-FLuc. After 16 hours, cells were placed in full medium for 48 hours before commencing antibiotic selection. To select stable cell lines, cells were grown in the presence of 500 µg/mL Geneticin (G418; Gibco) and 500 ng/mL Puromycin (Gibco).

### Protein Sample Preparation

Stably transfected cells were cultured to 70% confluency on a 100 mm dish, then washed twice using 10 mL phosphate buffered saline pH 7.4 (PBS, 137 mM NaCl/2.7 mM KCl/10 mM Na_2_HPO_4_). Four to five dishes of cells were then collected into a 1 mL lysis solution containing 850 μL radioimmunoprecipitation assay buffer (RIPA, 10 mM Tris-HCl pH 7.5/140 mM NaCl/1% NP-40/1% sodium deoxycholate/0.1% sodium dodecyl sulfate (SDS)) and 150 μL of Complete Mini protease inhibitor cocktail (Roche Diagnostic Systems, Laval, Canada). Harvested cells were sonicated using three 12-second bursts of a Sonic Dismembrator (model 500, Thermo Fischer Scientific, Ottawa, Canada) at an amplitude of 30%. Total amount of protein was quantified using the BCA assay (14).

### Western Blot

Protein samples of MDA-MB-435 cells stably expressing EGFP (40 µg), Tomato (40 µg), EGFP-MamE (40 µg), or Tomato-MamB (40 µg) were reduced with 100 mM dithiothreitol in sample preparation buffer (1 M Tris-HCl pH 6.8/10% SDS/0.1% Bromophenol Blue/43% glycerol) and heated at 85°C for at least 5 min. Reduced samples were then subjected to discontinuous SDS polyacrylamide gel electrophoresis (SDS-PAGE) using a 10% running gel. Protein was transferred onto a nitrocellulose blot using the Original iBlot Gel Transfer Device (Life Technologies, Burlington, Canada).

For EGFP detection, nonspecific protein binding was blocked in 5% bovine serum albumin (BSA)/Tris-buffered saline pH 7.4 (TBS) for 3 h at room temperature. Blots were then incubated for 15 h in 1:1000 mouse α-GFP (Invitrogen)/3% BSA/TBS/0.02 % sodium azide (TBSA); then washed using TBS/0.1% Tween 20 (TBST; Sigma-Aldrich, Oakville, Canada) for 30 min with 4 changes of buffer; and incubated for 2 h in 1:20,000 horseradish peroxidase (HRP)-conjugated goat α-mouse IgG (Sigma-Aldrich)/1% BSA/TBS. All incubations were performed at room temperature. Blots were then washed with 0.1% TBST for 30 min with 4 changes of buffer and imaged using the Chemigenius Gel Doc (Syngene). A chemiluminescent signal was detected using SuperSignal West Pico Chemiluminescent Substrate (Thermo Fischer Scientific), according to the manufacturer’s instructions.

For Tomato detection, blots were blocked in 3% BSA/TBSA for approximately 18 h at room temperature and then incubated for 18 h in 1:1000 primary goat α-tdTomato (MyBioSource, San Diego, USA)/3% BSA/TBSA at 4°C. After washing in 0.1% TBST as described above, blots were incubated for 1 h in 1:20,000 HRP-conjugated rabbit α-goat IgG (Sigma-Aldrich)/1% BSA/TBS at room temperature.

Glyceraldehyde 3-phosphate dehydrogenase (GAPDH) was used as a loading control. For GAPDH detection, blots were placed in stripping solution (1 M Tris-HCl pH 6.8/10% SDS/0.016% β-mercaptoethanol) and agitated in a 37°C water bath for 30 min prior to washing in 0.1% TBST and blocking in 5% BSA/TBS. The primary and secondary antibodies were 1:2000 rabbit α-GAPDH (Sigma-Aldrich)/3% BSA/TBSA and 1:20,000 HRP-conjugated goat α-rabbit IgG (Sigma-Aldrich)/1% BSA/TBS, respectively.

### Confocal Imaging

Stably transfected cell lines were examined with confocal fluorescence microscopy (using a Nikon A1R confocal microscope) to confirm expression and characterize the intracellular localization of EGFP-MamI, Tomato-MamL, Tomato-MamL_trunc,_ FLAG-MamL, EGFP-MamE (T317A), and Tomato-MamB fusion proteins. In preparation for confocal microscopy, approximately 100 thousand cells were cultured in a 35 mm glass-bottom dish (MatTek Corporation, Cedarlane, Burlington, Canada) for 48 hours. On the day of imaging, the dish was placed in a stage-top incubator to maintain 37°C and 5% CO_2_. Images and cines were captured using a Galvano scanner with NIS-Elements AR 5.11.01 (Nikon Instruments Inc.), and a 20x air objective with 0.75 numerical aperture. To capture images of cells expressing a single fluorophore, the FITC microscope filter (495 nm excitation/519 nm emission) was used for cells expressing the EGFP fluorophore and the TRITC microscope filter (557 nm excitation/576 nm emission) was used for cells expressing the Tomato fluorophore. To capture images of cells co-expressing both fluorophores (EGFP and Tomato), the FITC and TRITC filters allowing simultaneous 495 nm and 557 nm excitations was used. When needed, captured images of cells in both channels were then merged in Adobe Photoshop CS7.

Time-lapses were acquired with the time lapse function in NIS-Elements AR 5.11.01, recording an image every 1 s for a total of 60 s. Cines were captured in either channel or both channels simultaneously, as described above. The NIS-Elements software automatically generated a time lapse video in nd2 format with single or merged channels. This video was then edited in Adobe Photoshop CS7 by cropping and then pasting into a single file which could be exported in a graphics interchange format (GIF).

### Particle Trajectory Analysis

#### Particle Tracking

Using a plugin called Mosaic Particle Tracker 2D/3D (version 1.0.1) (15) in the software ImageJ version 1.8.0 (16), the trajectories of particles observed in cells expressing either Tomato-MamL, Tomato-MamL/EGFP-MamI, Tomato-MamL_trunc_, Tomato-MamB, or EGFP-MamE (T317A) were determined from confocal cines, and analyzed.

Cines converted to GIF files were loaded into ImageJ and prepared for analysis by converting them to greyscale; cropping to reduce their size and retaining only the portion of the movie with a single cell; then optimizing brightness and contrast. This last step does not affect the particle tracking but makes it easier for the user to visualize trajectories. For accurate estimation of diffusion coefficients and velocities, the pixel width d = L/N (where L is the width of the field of view in micron and N is number of pixels) and time interval between two consecutive frames τ = T/(F-1) (where T is the total duration of the video and F is the total number of frames) were calculated and added to the image properties in ImageJ.

After launching Mosaic Particle Tracker, the parameters used for particle detection were optimized for each cine: values of the radius (size of the tracked particles), cutoff (threshold intensity value below which detected particles are rejected), and percentile (range of intensities below the maximum intensity in the image for which fluorescent spots are considered to be particles) were manually adjusted to allow the software to detect the most manifest particles in the first image of the movie while not picking up lower-intensity noise speckle. For particle linking properties, the link range was set to 3 for all cines. In this way, the software would stop tracking a specific particle if it was absent for 3 consecutive frames. The displacement (maximum displacement allowed for a particle between two consecutive frames), which should be set to at least twice the average displacement of a particle between two frames, was set to 10 pixels. The software provided the total number of trajectories detected, a file containing the information relative to each detected trajectory (i.e., the position of the particle in each frame for which it was detected) and the MSD, <r^2^(t)>, of each detected particle.

### Simple Trajectory Analysis

The mobility of each particle was first assessed by fitting the MSD for each trajectory (within Mosaic Particle Tracker) with a simple power-law function:

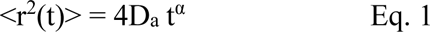

A particle undergoing free Brownian motion should have an MSD close to linear in time - that is, with an exponent α close to 1. Constrained Brownian motion would result in α < 1. In contrast, a particle undergoing directed motion should be characterized by α close to 2. If the motion is mixed, with alternate periods of Brownian and directed motion, then one would expect α to be somewhere between 1 and 2. As explained below, results of the fit of MSD was among the information used to differentiate between direct and Brownian motion. For the first group of particles, D_a_ can be considered to be an apparent diffusion coefficient (note that when α ≠ 1, this quantity does not have the dimension of a diffusion coefficient, as it only represents the diffusion coefficient that would be estimated from the fitted value of the MSD at t = 1 s, which is why it is more accurate to speak of an apparent diffusion coefficient).

### Refined Algorithm for Trajectory Analysis

Starting from the position and intensity of the tracked particle in each frame returned by Mosaic Particle Tracker, trajectories were further categorized and analyzed using an algorithm written for Mathematica. The main steps of this algorithm are as follows.

Trajectories were first evaluated for inclusion or exclusion based on the following criteria. Trajectories were rejected when either too short (less than 8 frames) or having a frame-to-frame displacement that was too large: that is, more than about 1 micron between successive frames or more than about 3 pixels between non successive frames, if the particle was not detected for one frame (these numbers were adjusted for each cell depending on the imaging parameters). Trajectories were also rejected if the average intensity of the particle was too low (less than one standard deviation below the mean intensity for all particles) or was detected for less than 70% of the trajectory duration.

For each of the remaining trajectories, the MSD was calculated and fitted (for lag times between 1 and 10 s) using Eq. 1. The velocity autocorrelation function (VAF), <v(0)v(t)> was also calculated for each of the remaining trajectories, in order to detect potential persistence in the direction of motion (examples of VAF are shown in the Supplementary data). An estimate of the particle maximum velocity was obtained by identifying the maximum value of the correlation between two successive measurements of the particle apparent velocity:

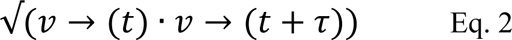

in which τ is the time interval between consecutive frames.

For all trajectories in a given cell, the distribution of single-step displacements (i.e., displacements between two successive frames, corresponding to a time interval τ = 1 s) was generated and fitted with the expression below, which accounts for the existence of two diffusive populations (i.e., the sum of two Rayleigh’s distributions):

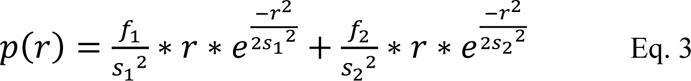

In some cases, a third term (a Gaussian peak attributed to steps taken during directed motion) was needed in order to account for a small fraction of longer displacement. Each of these populations is characterized by an average step size. The values of s_1_ and s_2_ (in order of increasing length) became the basis for sorting each individual trajectory into either an immobile, directed, or diffusive trajectory. The criteria used were as follows. For immobile trajectories, the total displacement and largest step are both less than 12 × s_1_. For a directed trajectory, the total displacement is more than 1.5 × s_2_ × √(*n*), where n is the number of steps in the trajectory, or the trajectory MSD is characterized by an exponent α > 1.1, as explained above in the section on simple trajectory analysis. Diffusive trajectories are those that are neither immobile nor directed.

### Statistical Analysis

All statistical tests were performed using GraphPad Prism version 8. A one-way analysis of variance (ANOVA) with a Tukey’s post hoc test was performed to determine any statistical significance between the apparent diffusion coefficient, velocity, and anomalous exponent values of the trajectories of magnetosome expression systems.

## Results

The following subsections describe the intracellular localization of essential magnetosome proteins as punctate structures, when expressed as fluorescent fusion proteins in a human melanoma cell line. The intracellular mobility of these structures is captured by confocal microscopy in movies and characterized according to particle trajectory, firstly using ImageJ software and a simple power law function, and latterly refined using a custom Mathematica algorithm.

### Cellular distribution of MamE and MamB

The magnetosome protein MamE consists of both conserved and variable sequences. We compared the expression of two EGFP-MamE fusion proteins, representing 3 distinct point mutations relative to the consensus sequence (Supplementary Figure S1), to examine potential differences in their intracellular expression in MDA-MB-435 cells. Using confocal fluorescence microscopy, cells stably expressing EGFP-*mamE* (G49S, T641S) display intracellular fluorescence in a punctate pattern with a diffuse fluorescence background (Figure 1A). These EGFP-MamE structures are numerous and cluster near the nucleus. Protein expression was confirmed by immunoblot (Figure 1B and Supplementary Figure S2) and reveals potential autoproteolysis of MamE as previously reported in MTB (12). Unlike the expression of MamL (10) or MamB (described below), MamE-expressing particles do not display mobility.

**Figure 1.**
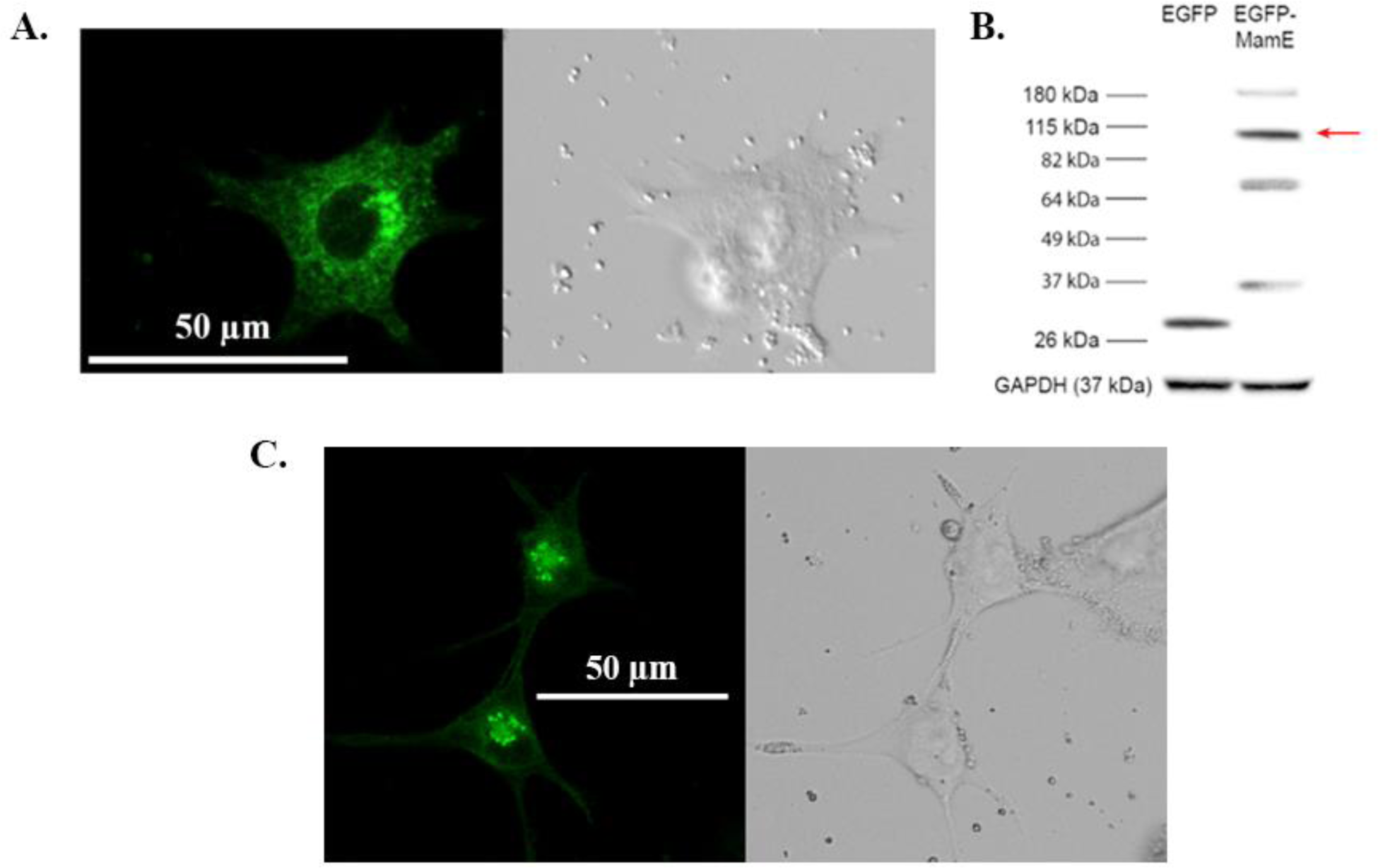
Expression of EGFP-MamE fusion proteins in mammalian cells. **A.** Confocal fluorescence microscopy of MDA-MB-435 cells stably expressing EGFP-*mamE* (G49S, T641S) reveals punctate intracellular fluorescence that clusters in the perinuclear region. **B.** Total cellular protein from cells stably expressing either EGFP or EGFP-*mamE* (G49S, T641S) were examined by western blot using a mouse α-EGFP primary antibody. The approximate size of EGFP-MamE is 110 kDa (red arrow). Bands at lower molecular weights are consistent with MamE autoproteolysis (16). Approximate MW is shown in the left margin. The bottom panel shows the loading control, GAPDH. **C.** Confocal fluorescence microscopy of MDA-MB-435 cells stably expressing EGFP-*mamE* (T317A) also displays a punctate intracellular fluorescence pattern.

Cells expressing EGFP-*mamE* (T317A) confirm that both *mam*E constructs show similar expression patterns. Confocal fluorescence microscopy of mammalian cells expressing EGFP-*mamE* (T317A) also shows distinct intracellular, punctate green fluorescence that clusters near the nucleus (Figure 1C). Similar to EGFP-MamE (G49S, T641S), the expression of EGFP-MamE (T317A) also exhibits a diffuse background and little or no mobility.

When Tomato-MamB is stably expressed in MDA-MB-435 cells, red fluorescence is displayed in a punctate pattern in the majority of the transfected cell population (∼70%; Fig. 2A) and in a diffuse pattern in about 30% of the transfected cell population (Fig. 2B). Importantly, these punctate MamB structures are mobile. Protein expression was confirmed by immunoblot (Figure 2C and Supplementary Figure S3).

**Figure 2.**
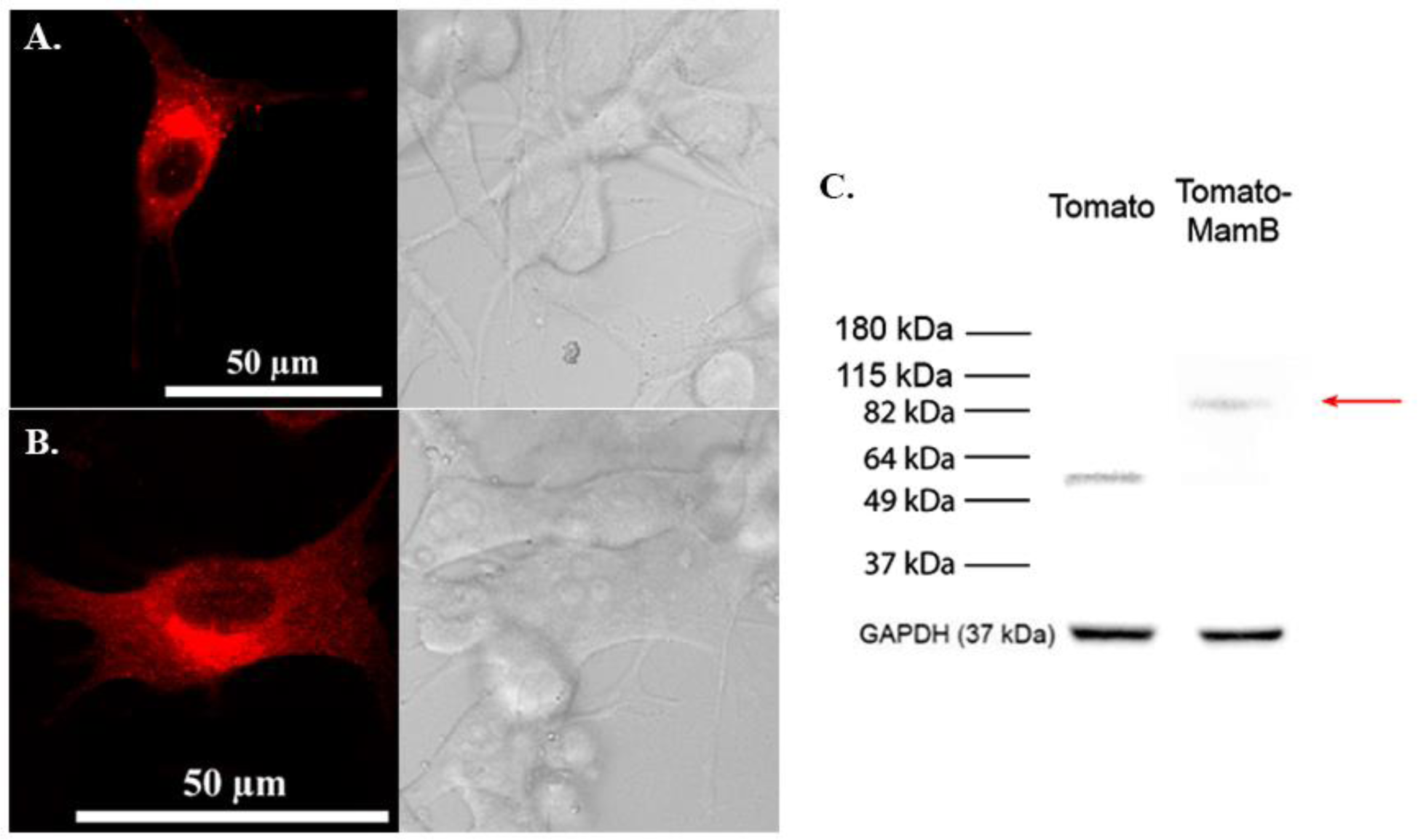
Expression of Tomato-MamB fusion protein in mammalian cells. Confocal fluorescence microscopy of MDA-MB-435 cells stably expressing Tomato-*mamB* displays either punctate intracellular fluorescence **(A)** or diffuse fluorescence **(B)**. The punctate structures in **(A)** are dispersed throughout the cell. Total cellular protein from cells stably expressing either Tomato or Tomato-MamB were examined by western blot **(C)** using a rabbit α-Tomato primary antibody. The approximate size of Tomato-MamB is 91 kDa (red arrow). Approximate MW is shown in the left margin. The bottom panel shows the loading control, GAPDH.

### Analysis of EGFP-MamE trajectories

As no evident motion of the puncta associated with MamE protein was observed in confocal movies, the analysis of EGFP-*mamE* (T317A)-expressing cells were chosen to serve as an example of cells containing particles with low mobility. Using this reference point, the signature of these MamE particles is contrasted with that of the higher mobility particles observed in other expression systems. A visual representation of MamE trajectories and the distribution of their displacement is shown in Figure 3 (and Supplementary Figure S4). It is apparent from this distribution that the motion of MamE particles is very highly constrained, since almost no single displacement after t = 1s is larger than 0.2 µm, and since the displacements after 20 s are almost the same as after 1 s. Trajectory analysis of MamE fluorescent structures showed that 26% of these particles were immobile while 62% were undergoing Brownian motion and only 13% underwent directed motion at some point in their trajectories (Table 2). According to this analysis, the average apparent diffusion coefficient of MamE particles undergoing Brownian motion is 1.9 ± 0.4 x 10^-3^ µm^2^/s. The average velocity of the very few MamE particles undergoing directed motion is 0.17 ± 0.03 µm/s.

**Figure 3.**
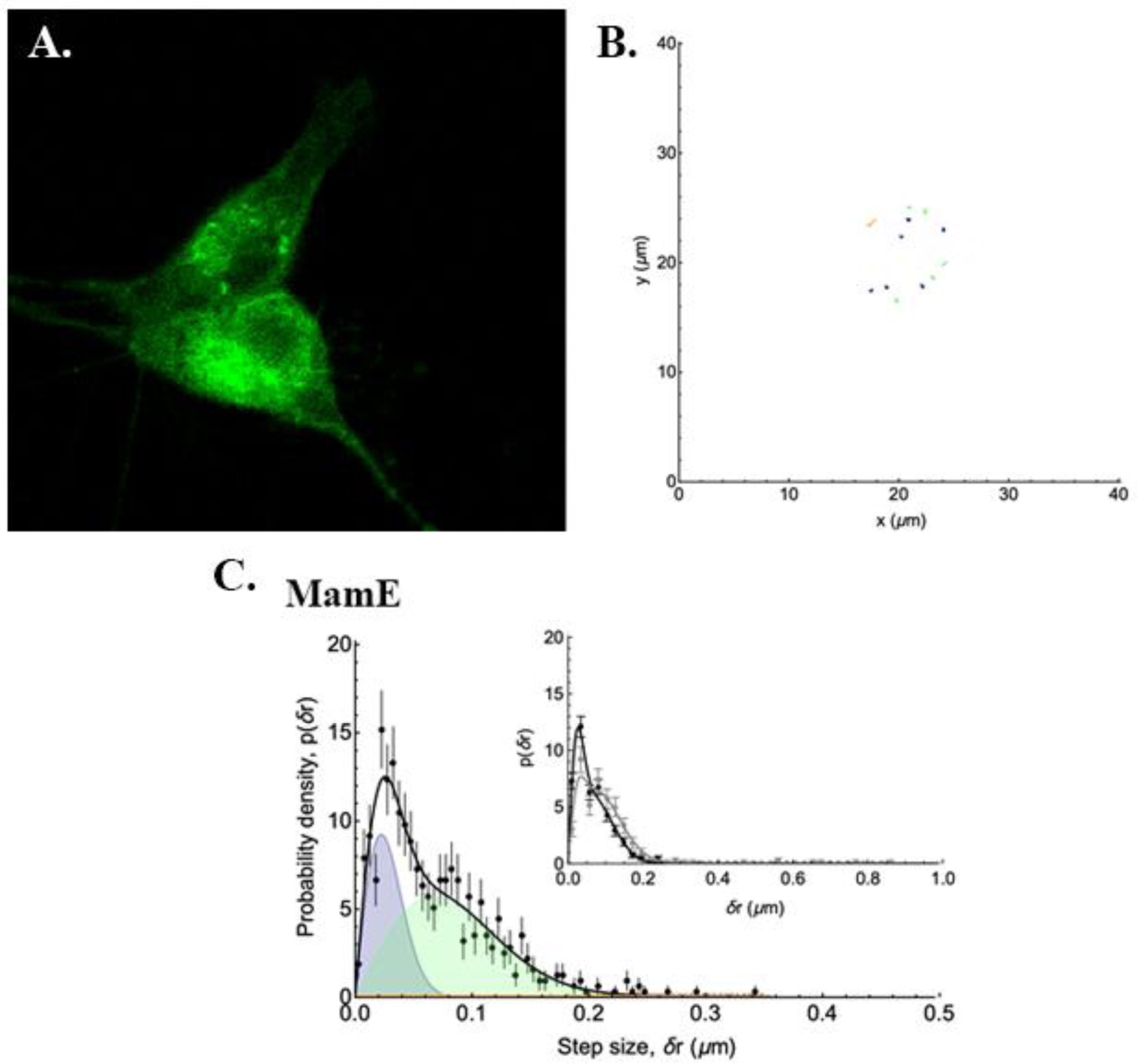
Distribution of the displacements of punctate structures in a representative EGFP-MamE-expressing mammalian cell. The analysis of punctate structures in the confocal micrograph of a representative MamE-expressing cell **(A)** reveals 3 types of trajectories **(B)** with stationary particles shown in dark blue, and Brownian motion and directed motion trajectories represented in green and orange, respectively. The probability density of displacements **(C)** after t = 1s is shown in the main panel (black symbols) and fitted assuming two diffusive populations and one directed population. The blue and green shaded peaks represent the contributions of the immobile and Brownian diffusing populations, the orange shaded peak - barely visible in this case – is that of the directed population and the black curve shows the sum of these three contributions. The inset shows very little difference in the distribution of displacements after 1 s (black symbols) and 20 s (grey symbols).

**Table 2.**
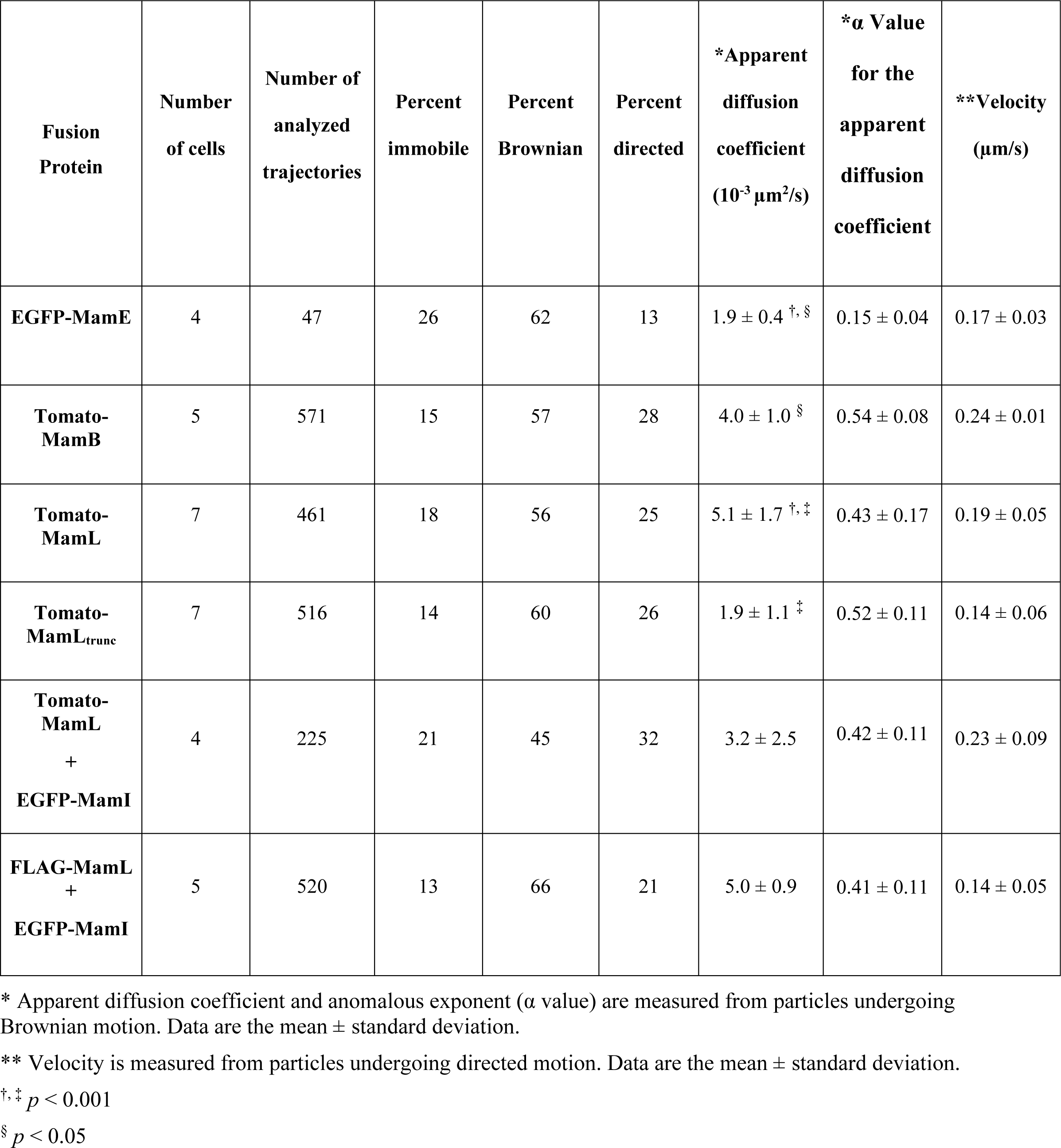
Summary of trajectory analysis in mammalian cells expressing magnetosome proteins.

### Mobility of Tomato-MamB

Analyses of Tomato-MamB particle trajectories also reveal both directed and Brownian motion in each cell, with the occurrence of directed trajectories markedly greater than in MamE-expressing cells. A visual representation of the detected MamB particles (Figure 4A) and analyzed trajectories (Figure 4B) is provided alongside their displacement distribution (Figure 4C and Supplementary Figure S5). The difference between MamE and MamB is clearly shown by the large number of displacements after t = 1 s between 0.2-0.4 µm and by the broad distribution of displacements after 20 s (Figure 4C inset), as expected for mobile particles.

**Figure 4.**
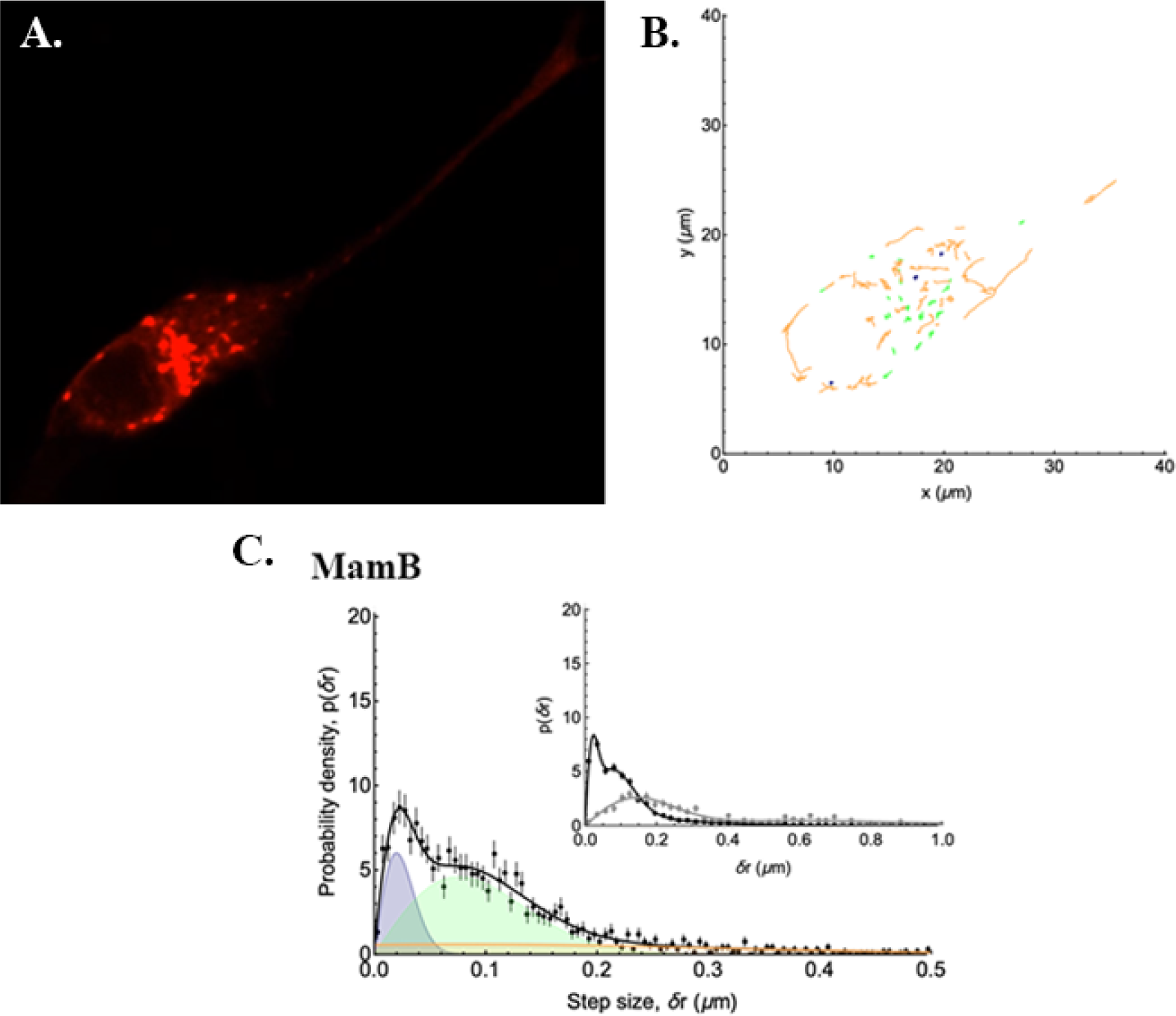
Distribution of the displacements of punctate structures in a representative Tomato-MamB-expressing mammalian cell. The analysis of punctate structures in the confocal micrograph of a representative MamB-expressing cell **(A)** reveals 3 types of trajectories **(B)** with stationary particles shown in dark blue, and Brownian motion and directed motion trajectories represented in green and orange, respectively. The probability density of displacements **(C)** after t = 1s is shown in the main panel (black symbols) and fitted assuming two diffusive populations and one directed population. The blue and green shaded peaks represent the contributions of the slow and fast diffusing populations, the orange shaded peak that of the directed population and the black curve shows the sum of these three contributions. The inset shows a broad distribution of displacements after t = 20 s (grey symbols) compared to t = 1 s (black symbols).

Analysis of MamB trajectories showed that only 15% of MamB particles were immobile, while 57% were undergoing Brownian motion, and 28% underwent directed motion at some point in their trajectories (Table 2). The average apparent diffusion coefficient of MamB particles undergoing Brownian motion is 4.0 ± 1.0 x 10^-3^ µm^2^/s and 2-fold greater than MamE particles (p < 0.05). The average velocity of MamB particles undergoing directed motion is 0.24 ± 0.01 µm/s.

### Mobility of Tomato-MamL

Previous characterization of Tomato-MamL particles (10) revealed patterns of mobility similar to those of Tomato-MamB. A visual representation of detected Tomato-MamL particles and analyzed trajectories is shown in Figures 5A and 5B, respectively. The distribution of their displacements is plotted in Figure 5C (and Supplementary Figure S6), with a broad distribution apparent after 20 s. Analysis of these results showed that 18% of MamL particles were immobile; 56% were undergoing Brownian motion; and 25% underwent directed motion at some point in their trajectories (Table 2) as seen for MamB-expressing particles. The average apparent diffusion coefficient of MamL particles undergoing Brownian motion is 5.1 ± 1.7 x 10^-3^ µm^2^/s. The average velocity of MamL particles undergoing directed motion is 0.19 ± 0.05 µm/s. Similar to Tomato-MamB, the apparent diffusion coefficient of Tomato-MamL particles is over 6-fold greater than EGFP-MamE (p < 0.001, Table 2) while the velocity of any one particle remains approximately the same between expression systems (Table 2).

**Figure 5.**
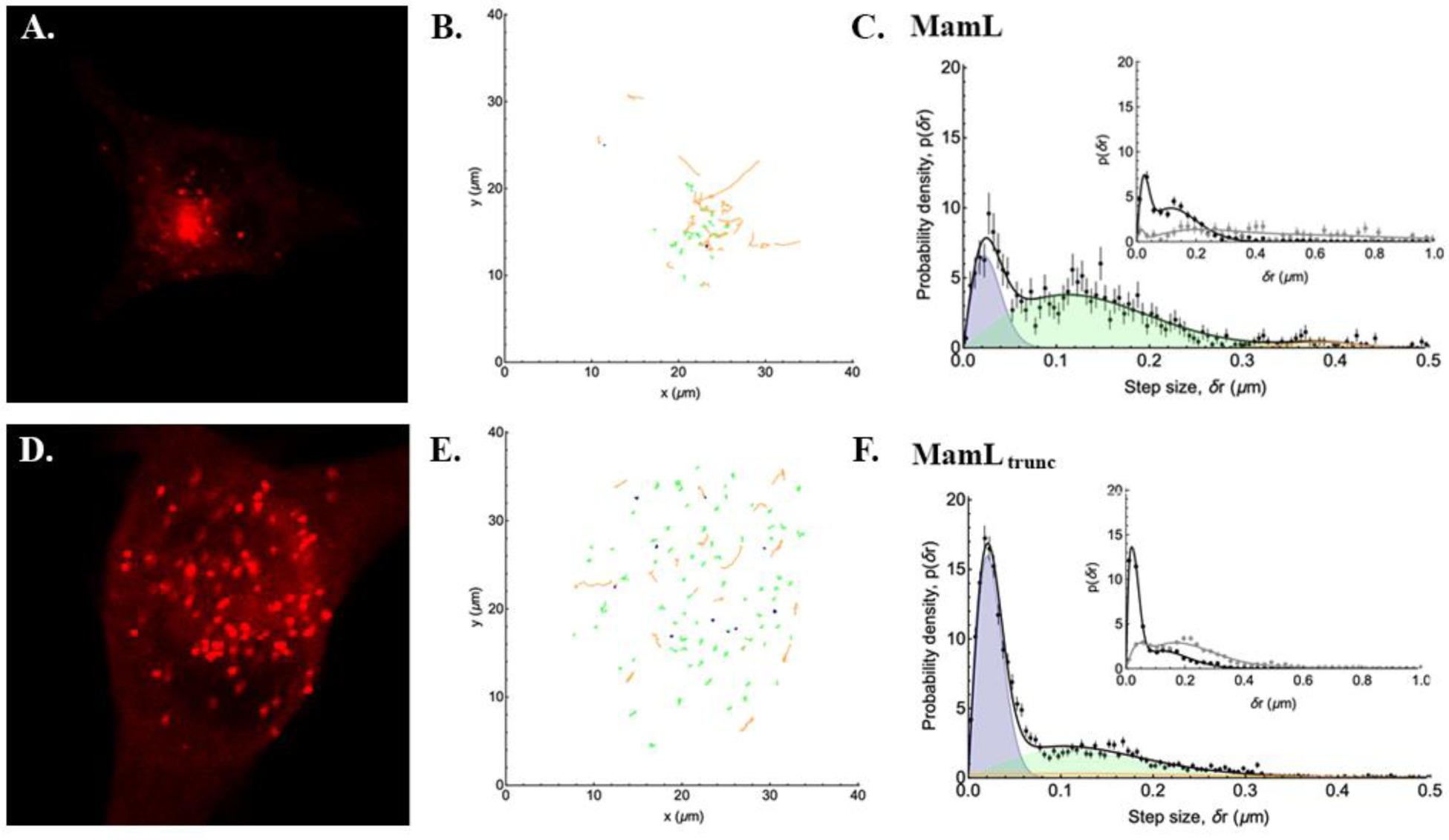
Comparing the displacement distributions of punctate structures in representative MamL- and MamL_trunc_-expressing mammalian cells. The analysis of punctate structures in the confocal micrograph of a representative MamL-expressing cell **(A)** reveals 3 types of trajectories **(B)** with stationary particles shown in dark blue, and Brownian motion and directed motion trajectories represented in green and orange, respectively. The probability density of displacements **(C)** after t = 1 s is shown in the main panel (black symbols) and fitted assuming two diffusive populations and one directed population. The blue and green shaded peaks represent the contributions of the slow and fast diffusing populations, the orange shaded peak that of the directed population and the black curve shows the sum of these three contributions. The inset shows a broad distribution of displacements after t = 20 s (grey symbols) compared to t = 1 s (black symbols). These data are compared to a representative MamL_trunc_-expressing cell **(D)** and the displacement of its punctate structures **(E)**. The probability density of displacements **(F)** plotted after t = 20 s (inset) shows a persistent broad distribution despite removal of the MamL C-terminal peptide.

To better understand the mobility of MamL, we introduced an early stop codon to remove 15 amino acids from the C terminus of MamL protein and examined this truncated form. The C-terminal peptide that was removed may interact with mobile cytoskeletal elements based on sequence similarity to cell-penetrating peptides (2). A visual representation of the detected MamL_trunc_ particles and analyzed trajectories is shown in Figures 5D and 5E, respectively, with their displacement distribution plotted in Figure 5F (and Supplementary Figure S7). Despite truncation of its C-terminal peptide, there remains a broad distribution of displacements after t = 20 s (inset). Analysis of MamL_trunc_ trajectories showed that 14% of MamL_trunc_ particles were immobile, 60% were undergoing Brownian motion, and 26% underwent directed motion at some point in their trajectories (Table 2). While the distribution of particles is similar to native MamL, the average apparent diffusion coefficient of MamL_trunc_ particles undergoing Brownian motion was reduced more than 2.5-fold (p < 0.001) to 1.9 ± 1.1 x 10^-3^ µm^2^/s (Table 2). The average velocity of MamL_trunc_ particles undergoing directed motion is 0.14 ± 0.06 µm/s and comparable to all other particles expressing magnetosome proteins (Table 2).

### Mobility of co-expressed Tomato-MamL/EGFP-MamI

Tomato-MamL particles not only retain their mobility when co-expressed with EGFP-MamI, the latter appear to be recruited to the same mobile complex (10). These mobile particles, consisting of co-localized protein, also display both directed and Brownian motion. A visual representation of the detected MamL+MamI particles and trajectories is shown in Figures 6A and 6B, respectively, with their displacement distribution plotted in Figure 6C (and Supplementary Figure S8). Similar to the mobility of particles in other MamL expression systems, there is a broad distribution after t = 20 s (inset). Analysis of the MamL+MamI trajectories revealed that 21% of MamL+MamI particles were immobile, 45% were undergoing Brownian motion, and 32% underwent directed motion at some point in their trajectories (Table 2). Compared to particles expressing MamE, these data suggest that co-expression of essential magnetosome proteins more than doubles the fraction of particles that undergo directed motion. The average apparent diffusion coefficient of MamL+MamI particles undergoing Brownian motion is 3.2 ± 2.5 x 10^-3^ µm^2^/s. The average velocity of MamL+MamI particles undergoing directed motion is 0.23 ± 0.09 µm/s, consistent with the other expression systems examined (Table 2).

**Figure 6.**
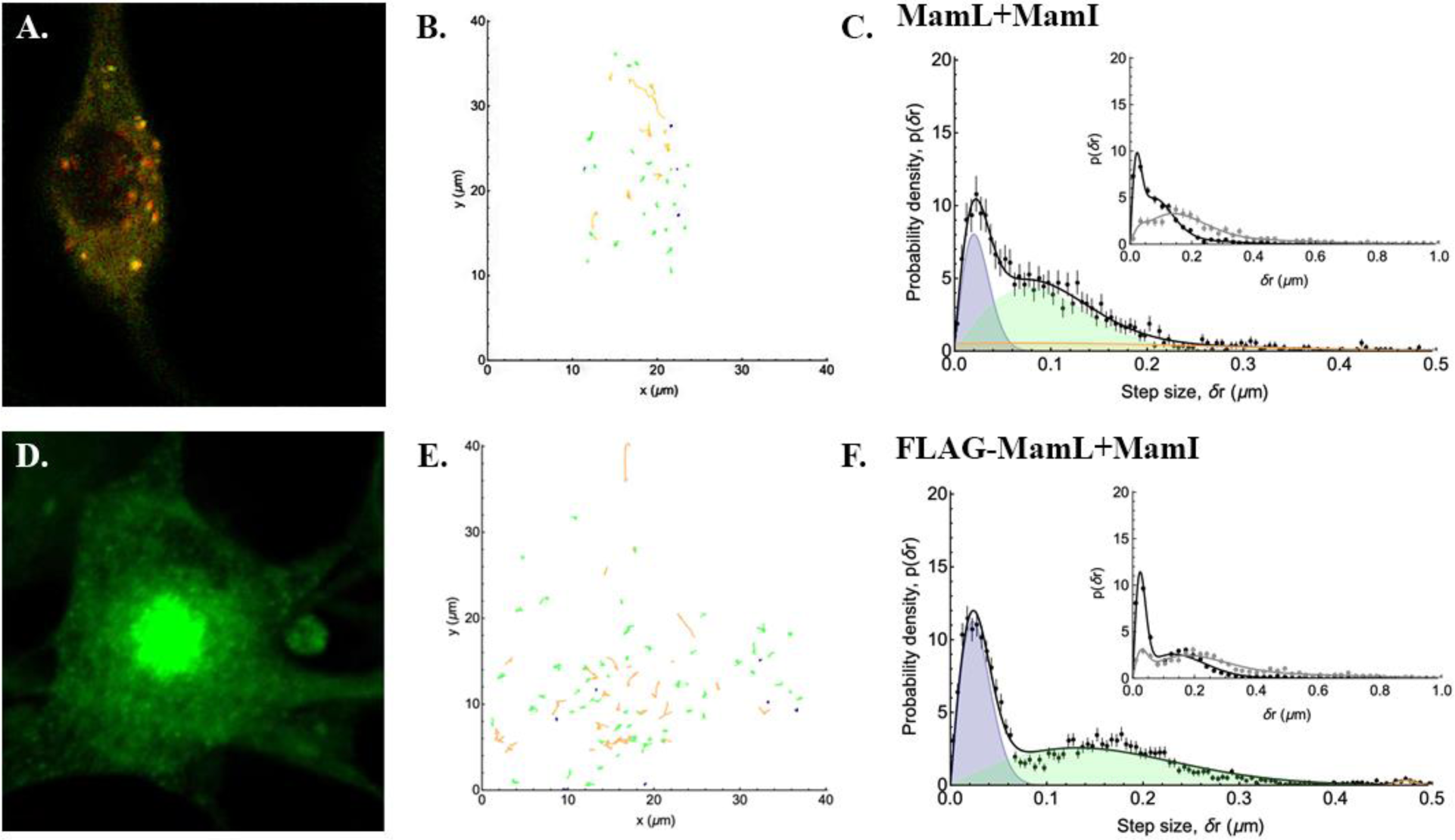
Comparing the displacement distributions of punctate structures in representative mammalian cells co-expressing MamL+MamI with or without Tomato fluorescence. The analysis of punctate structures in the confocal micrograph of a representative cell co-expressing Tomato-MamL+EGFP-MamI (**A**, yellow fluorescence) reveals 3 types of trajectories **(B)** with stationary particles shown in dark blue, and Brownian motion and directed motion trajectories represented in green and orange, respectively. The probability density of displacements **(C)** after t = 1 s is shown in the main panel (black symbols) and fitted assuming two diffusive populations and one directed population. The blue and green shaded peaks represent the contributions of the slow and fast diffusing populations, the orange shaded peak that of the directed population, and the black curve shows the sum of these three contributions. The inset shows a broad distribution of displacements after t = 20 s (grey symbols) compared to t = 1 s (black symbols). By replacing Tomato with a FLAG sequence, a comparison is drawn to representative data from FLAG-MamL+EGFP-MamI co-expression (**D**, green fluorescence) and the displacement of these punctate structures **(E)**, with the probability density of displacements **(F)** plotted after t = 1 s and t = 20 s (inset) broadening with time.

A FLAG-MamL/EGFP-MamI co-expression system was created to simplify confocal image analysis using a single chromophore (EGFP) and to remove bulky fluorescent proteins attached to MamL (i.e., tandem Tomato) that may compromise magnetosome protein interactions (10). The trajectories of particles from stable FLAG-MamL/EGFP-MamI co-expression were analyzed and compared to those of the dual fluorescent Tomato-MamL/EGFP-MamI particles. A visual representation of detected FLAG-MamL+MamI particles and their displacements is shown in Figures 6D and 6E, respectively, with the displacement distribution plotted in Figure 6F (and Supplementary Figure S9). Similar to Tomato-MamL/EGFP-MamI, the analysis of FLAG-MamL/EGFP-MamI trajectories showed that a minority of the particles were immobile (13%) while the majority of particles were undergoing Brownian motion (66%) and directed motion (21%) at some point in their trajectories (Table 2). The average apparent diffusion coefficient of FLAG-MamL+MamI particles undergoing Brownian motion is 5.0 ± 0.9 x 10^-3^ µm^2^/s (Table 2). The average velocity of FLAG-MamL+MamI particles undergoing directed motion is 0.14 ± 0.05 µm/s and once again comparable to the other particles formed by the essential magnetosome proteins examined (Table 2).

## Discussion

This is the first report to describe the stable expression of MamE and MamB in mammalian cells and analyze the intracellular mobility of punctate structures formed by essential magnetosome proteins expressed in this eukaryotic environment. For each fluorescent fusion protein analyzed (MamL, MamL_trunc_, MamL+MamI, MamB, or MamE), three types of particle movement were observed for the puncta: confined trajectories, Brownian trajectories, and trajectories with stretches of directed motion.

Both MamE and MamB fusion proteins displayed intracellular, punctate fluorescence similar to the pattern of MamL expression (10). However, EGFP-MamE-expressing cells often displayed punctate fluorescence in the perinuclear region while approximately 30% of Tomato-MamB-expressing cells had no discernable punctate fluorescence. Also unlike MamL expression, both MamB- and MamE-expressing cells displayed diffuse fluorescence.

Trajectories undergoing Brownian motion have apparent diffusion coefficients between 2 x 10^-3^ and 5 x 10^-3^ µm^2^/s. This diffusion coefficient is very slow compared to that of a soluble protein in the cytoplasm of mammalian cells, which is on the order of 10 µm^2^/s (17). Assuming that MamI, MamL, MamB, and MamE are all membrane associated proteins when expressed in mammalian cells, as in MTB, the measured diffusion coefficients are consistent with the notion that these magnetosome proteins are part of a rather large (membranous) structure and not simply part of the soluble protein fraction. Since these punctate structures cannot be resolved by confocal microscopy, the radius of these fluorescent particles is expected to be approximately less than or equal to 100 nm (i.e., smaller than the resolution of the microscope). Hence, the possibilities for particles of this diameter include localization in the membrane of pre-existing lipid vesicles (18) or a role in the formation of new ones. Confined trajectories may thus correspond to particles localized in immobilized vesicles.

Particles that undergo active directed motion have a maximum velocity from 0.1 to 0.3 µm/s. These particles usually also undergo stretches of Brownian motion or immobility. The fluorescent magnetosome fusion proteins may be localized in vesicles that are attached to molecular motors and/or may be directly interacting with molecular motors as they transiently walk along protein filaments.

### Brownian motion of magnetosome proteins

MamE particles are either immobile (26%) or undergoing some form of Brownian motion (62%); however, several features of this analysis point toward a restricted type of Brownian motion. First, Brownian MamE particles have an apparent diffusion coefficient of 1.9 ± 0.4 x 10^-3^ µm^2^/s, which is significantly lower than the diffusion of MamL and MamB particles (Table 2). This suggests that MamE is localized in different structures than MamL and MamB, with more restricted mobility. Additionally, Brownian MamE particles have trajectories characterized by an extremely low anomalous exponent of 0.15 ± 0.04, the lowest α value of all the magnetosome protein expression systems examined. An α value below 1 points to constrained or restricted diffusion (i.e. diffusion in the presence of obstacles), and a value below 0.5 further points towards caged diffusion (19, 20). Taken together, the low diffusion coefficient of MamE particles, extremely low value of the anomalous exponent α, and the high occurrence of stationary MamE particles suggest that this protein is the most constrained out of the magnetosome proteins studied in mammalian cells.

In contrast to MamE, Brownian MamL and MamB particles have a noticeably larger diffusion coefficient (on the order of 4-5 x 10^-3^ µm^2^/s), and a larger anomalous exponent α (on the order of 0.4 to 0.5). These values are consistent with magnetosome proteins localizing on particles (e.g. lipid vesicles) that are tens to hundreds of nm in size and undergoing restricted diffusion (18). MamL_trunc_ particles, which diffuse at 1.9 ± 1.1 x 10^-3^ µm²/s, diffuse significantly slower than MamL particles (*p* < 0.001). Removal of the MamL C-terminal peptide may cause the protein to localize in different (more confined) structures than full-length MamL. Alternatively, the MamL C-terminal may confer mobility, either through its association with the particle structure or by binding directly to molecular motors.

When MamL is co-expressed with MamI, the diffusion coefficient of Tomato-MamL/EGFP-MamI particles (3.2 ± 2.5 x 10^-3^ µm^2^/s) is no different than the diffusion of structures with MamL alone (5.1 ± 1.7 x 10^-3^ µm^2^/s). This is consistent with the conclusion that MamL and MamI are localized within the same structures/particles, and that MamI is recruited to the same structure as MamL when they interact (10). This is further supported by the FLAG-MamL/EGFP-MamI expression system. These particles diffuse at 5.0 ± 0.9 x 10^-3^ µm^2^/s, which is not significantly different from the diffusion coefficient of particles containing either Tomato-MamL/EGFP-MamI or Tomato-MamL alone. Furthermore, the α value of Tomato-MamL alone, Tomato-MamL/EGFP-MamI, and FLAG-MamL/EGFP-MamI particles are all comparable.

### Directed motion of magnetosome proteins

Although the magnetosome proteins studied undergo directed motion to a different extent (i.e. variability in the percentage of detected particles that display directed motion and in the average value of the α exponent for these directed trajectories; Supplementary Table S1), similar velocities (around 0.2 µm/s) were measured for all particles. In addition, except for MamE particles, the aspect of these directed trajectories (which very often are accompanied by a period of intermittent Brownian motion) is similar for all studied proteins (Figure 7).

**Figure 7.**
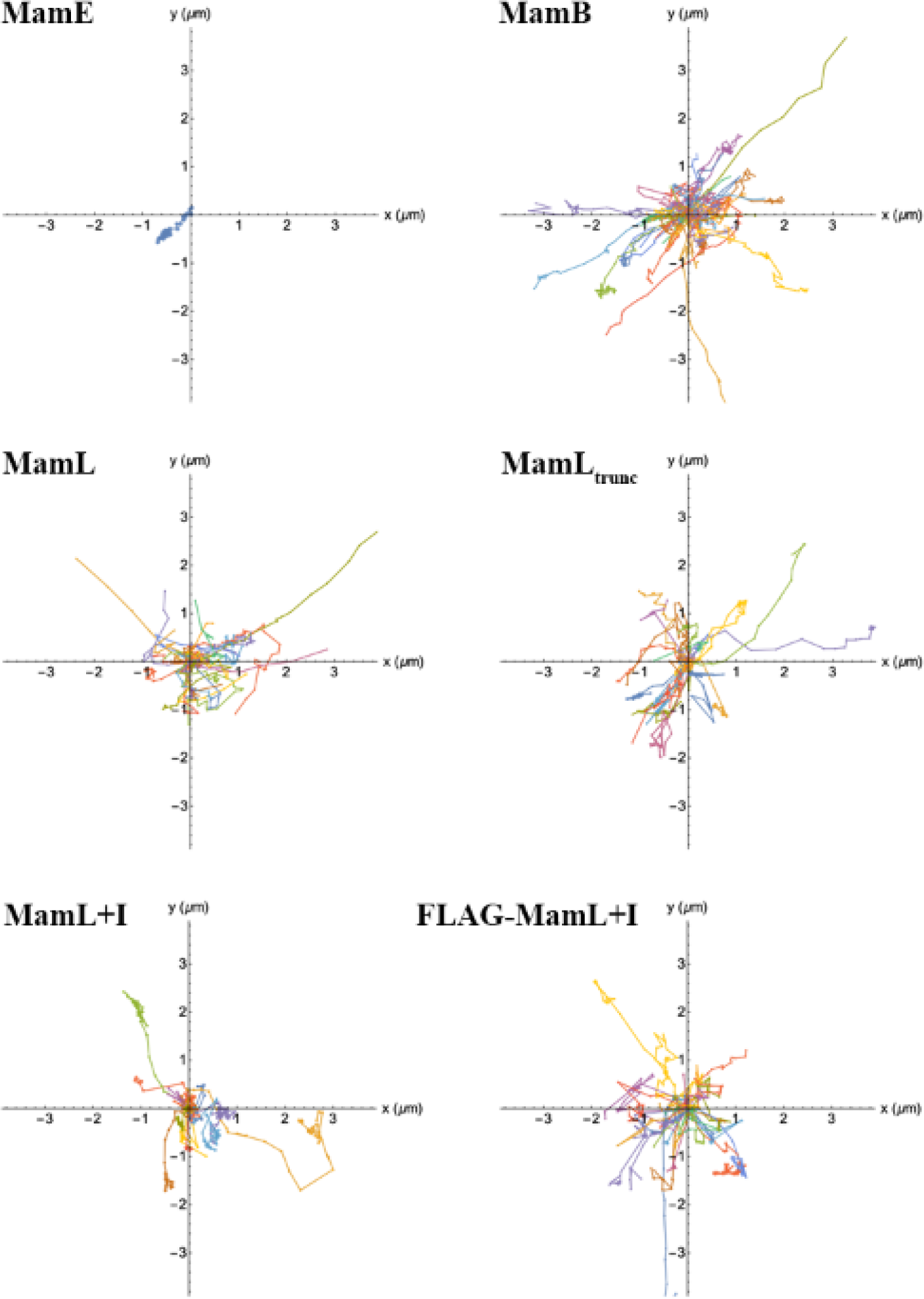
Representative directed trajectories of magnetosome proteins expressed in mammalian cells. Each panel shows all the directed trajectories detected in a single representative cell, for each of the different expression systems under study. All trajectories have been plotted by placing the first point of the trajectory at the point with coordinates x = y = 0. Each coloured line represents the distance and direction travelled by a single trajectory.

The directed motion observed suggests that detected particles interact with some kind of linear molecular motor present in the cell, for example a type of myosin motor. In mammalian cells, myosin motors are a group of molecular motor proteins responsible for directed movement of intracellular cargo. These structures, specifically myosin IB, myosin II, and myosin V, all move at a velocity of 0.2 µm/s (21). The speeds observed for the detected particles are therefore consistent with interactions with myosin. Molecular motors with a negative charge, or that associate with cargo adaptor proteins that are negatively charged, may interact with positively-charged proteins in the cell (11).

MamE stands out from other magnetosome proteins studied here in that many fewer MamE particles are undergoing directed motion and/or they spend less time undergoing active directed motion (Figure 7). This is further indicated by the low α exponent of 0.26 ± 0.32 (Supplementary Table S1). Particles undergoing directed motion should have α exponents close to 2; however, a lower exponent can also indicate that the directed motion alternates with either Brownian motion or immobilized motion, as is the case with all particles detected and especially with MamE. When undergoing directed motion, MamE particles move at a velocity of 0.17 µm/s. From protein structure analysis, the cytoplasmic C-terminal polypeptide of MamE is short and has only 2 positively-charged amino acids; therefore, MamE most likely does not bind to molecular motors strongly or frequently due to the small positive charge. Alternatively, MamE may be located in a vesicle that does not interact often with molecular motors. This may explain why MamE is not undergoing directed motion as often and why the protein seems mostly immobile while observed under a confocal fluorescence microscope.

MamB particles undergoing active directed motion travel at 0.24 µm/s. From protein structure analysis, the cytoplasmic domain of MamB is quite large and almost 20% of its amino acids have positive charges. These positive charges, therefore, have opportunities to interact with negatively-charged molecular motors in the mammalian cytoplasm. We can thus infer that MamB either interacts directly with molecular motors or is located in a vesicle that interacts with molecular motors.

MamL particles undergoing active directed motion move at approximately 0.2 µm/s, and are thus expected to also be interacting with molecular motors (i.e., myosin). Although the cationic amino acid residues of the MamB and MamL cytoplasmic domains are not arranged in a similar pattern, we can speculate that these two proteins may be interacting with similar molecular motors since they move at similar velocities. MamL_trunc_ particles that are undergoing active directed motion travel at 0.14 µm/s, which is not significantly different from the velocity of full-length MamL particles. It should be noted that the α exponent of MamL_trunc_ particles is smaller than that of MamL particles (0.88 ± 0.31 vs. 1.01 ± 0.18). This result aligns with what is seen with the confocal microscope: that MamL_trunc_ particles do not undergo directed motion as frequently as full-length MamL particles (10).

Tomato-MamL/EGFP-MamI and FLAG-MamL/EGFP-MamI particles undergoing active directed motion move at similar velocities as Tomato-MamL alone (0.23 µm/s vs. 0.14 µm/s vs. 0.20 µm/s, respectively). This suggests that Tomato-MamL alone, Tomato-MamL/EGFP-MamI, and FLAG-MamL/EGFP-MamI particles interact with the same molecular motors. Unlike the predicted protein structures of MamL, MamI has no cytoplasmic domain and thus has no opportunity to interact with molecular motors (22). Whether expressed alone or together with MamI, MamL is the protein that likely interacts with molecular motors.

## Conclusion

The mobility of magnetosome proteins MamL, MamB, and MamE in a mammalian system indicates their possible interaction with mammalian molecular motors. The pattern of motion of the structures to which these proteins localize is consistent with the motion of lipid vesicles (endosomes, lysosomes) observed in mammalian cells, namely a mixture of Brownian motion (with very small diffusion coefficient on the order of 10^-3^ to 10^-2^ µm^2^/s) and directed motion (with velocity on the order of 0.1 to 1 µm/s; Supplementary Figure S10) (23, 24).

Moreover MamL, MamB, and MamE all move at similar velocities when undergoing active directed motion. This suggests that, when expressed in mammalian cells, these magnetosome proteins may interact with the same type of molecular motor or localize to a structure that interacts with a common type of molecular motor.

When co-expressed, MamI and MamL localize to the same intracellular compartment and form punctate, mobile structures. Importantly, MamL+MamI particles undergoing directed motion move at the same velocity as particles in cells expressing MamL alone, suggesting they interact with the same molecular motor. In addition, MamL+MamI particles diffuse slower than particles with MamL alone, consistent with the expected slower diffusion of larger structures relative to smaller ones.

Taken together, this trajectory analysis supports mounting evidence that magnetosome proteins are compatible with mammalian cell systems, and localize and interact with mammalian membranous compartments. The diffusion coefficients and similar velocities at which these essential magnetosome proteins travel inside the cell are consistent with their proposed localization on an intracellular membrane. Finally, the evidence for co-localization and interaction of MamI and MamL is further substantiated through analysis and comparison of the mobility of particles with MamL alone versus MamL+MamI. This study is thus consistent with our hypothesis that essential magnetosome proteins MamI, MamL, MamB, and MamE interact in a membranous intracellular compartment in mammalian cells.

## Acknowledgements

The authors thank Moeiz Ahmed for preparing Supplementary Figure S1. This study was supported by Discovery Grants (2014–05589, 2020–04125) from the Natural Sciences and Engineering Research Council of Canada.

## Database

The Mathematica code described herein may be accessed at the following site: https://github.com/cecilefradin/Particle-Trajectory-Classification-and-Analysis/.

## Supplementary Tables and Figures

**Supplementary Figure S1.**
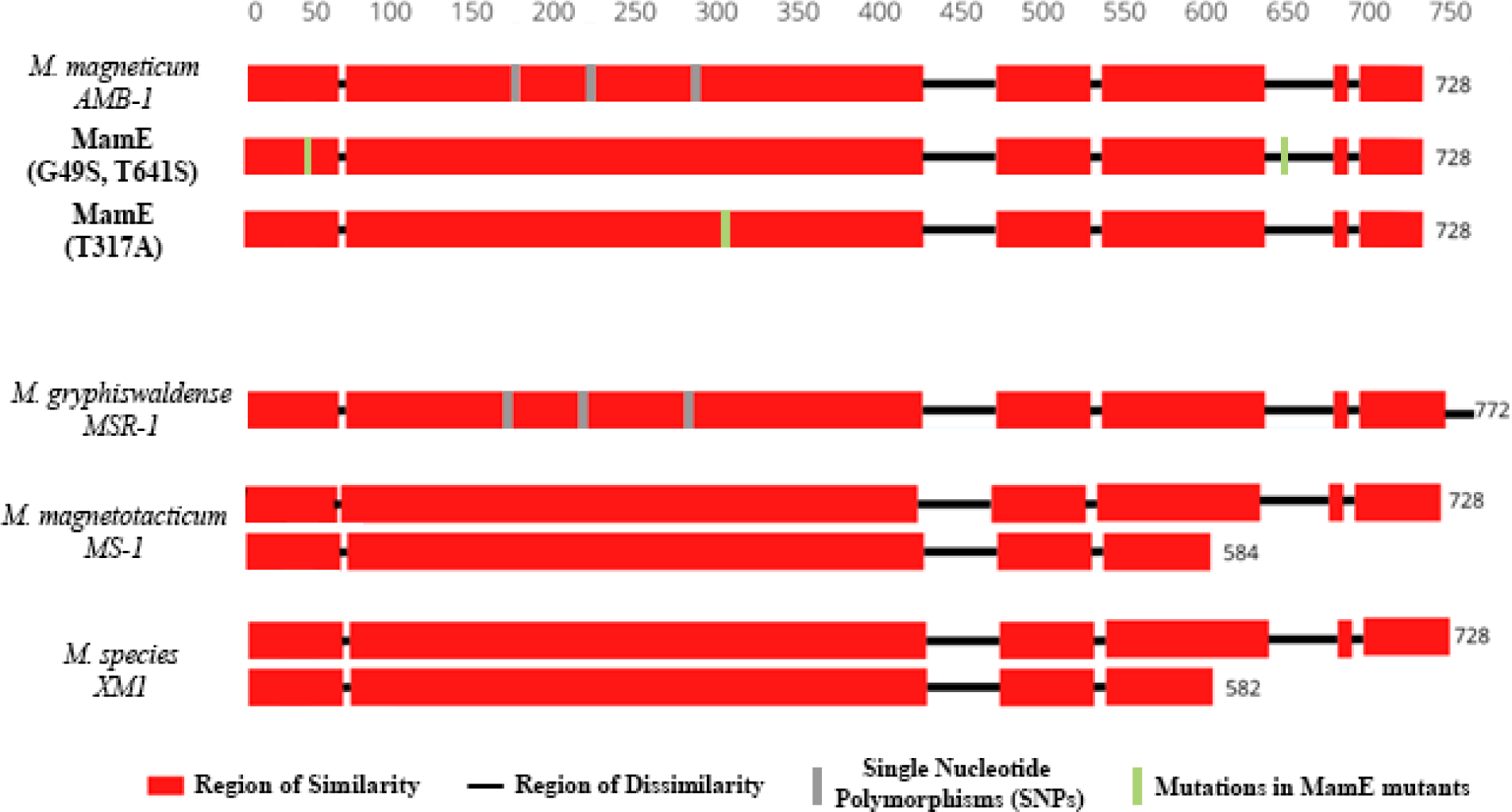
MamE sequence alignment and homology across several magnetotactic bacterial species. Regions of similarity in the amino acid sequence of MamE across species are highlighted in red, while regions of dissimilarity are in black, and single nucleotide polymorphisms (SNPs) appear in grey. Minor changes in the sequence of MamE (G49S, T641S) and MamE (T317A) are indicated in green and compared to the sequence from *Magnetospirillum magneticum* AMB-1 from which they were cloned. While the conservative substitution (T641S) in a region of variability is not expected to influence MamE function, neither substitution in different conserved regions (G49S or T317A) altered the trajectory analysis of MamE expressed in mammalian cells.

**Supplementary Figure S2.**
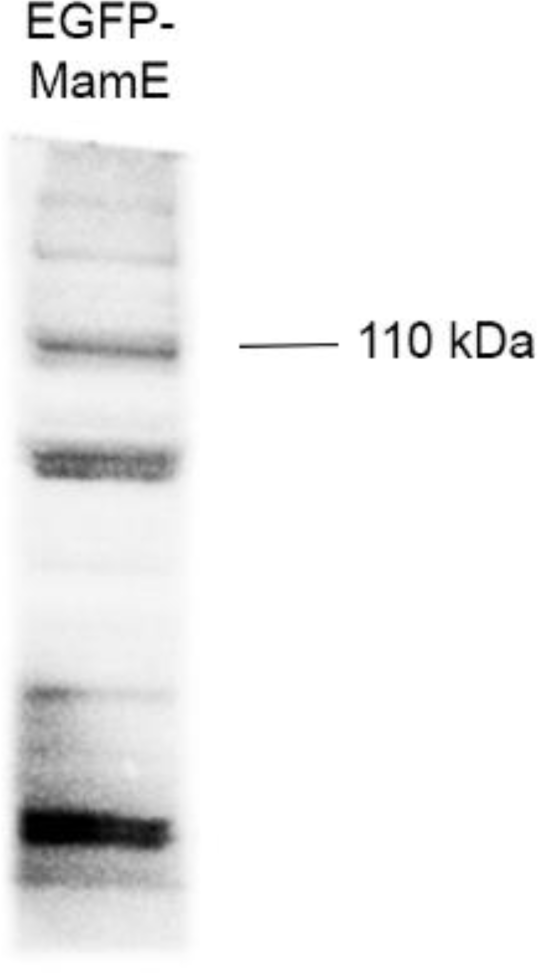
Immunoblot of total cellular protein from mammalian cells expressing EGFP-MamE. Total cellular protein from MDA-MB-435 cells stably expressing EGFP-MamE was examined with mouse α-EGFP. The full-length western blot is displayed.

**Supplementary Figure S3.**
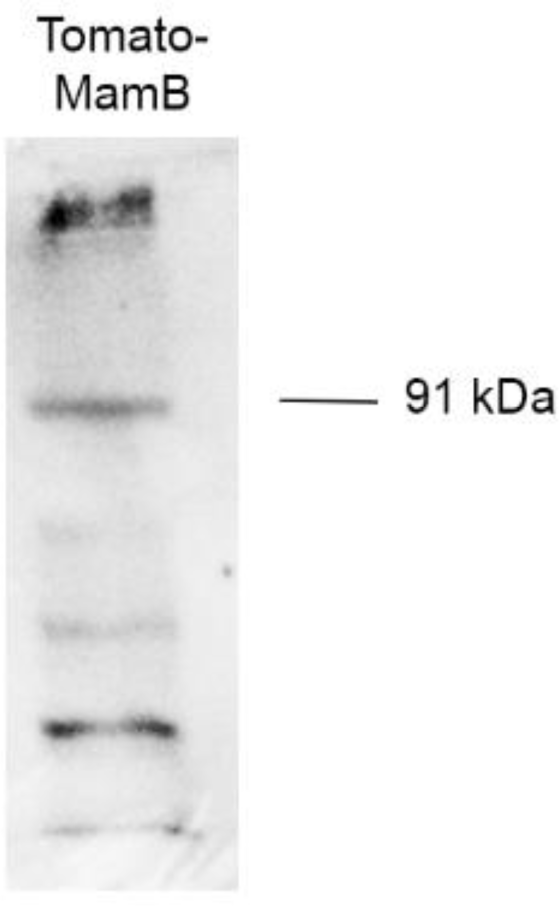
Immunoblot of total cellular protein from mammalian cells expressing Tomato-MamB. Total cellular protein from MDA-MB-435 cells stably expressing Tomato-MamB was examined with rabbit α-Tomato. The full-length western blot is displayed.

**Supplementary Figure S4:**
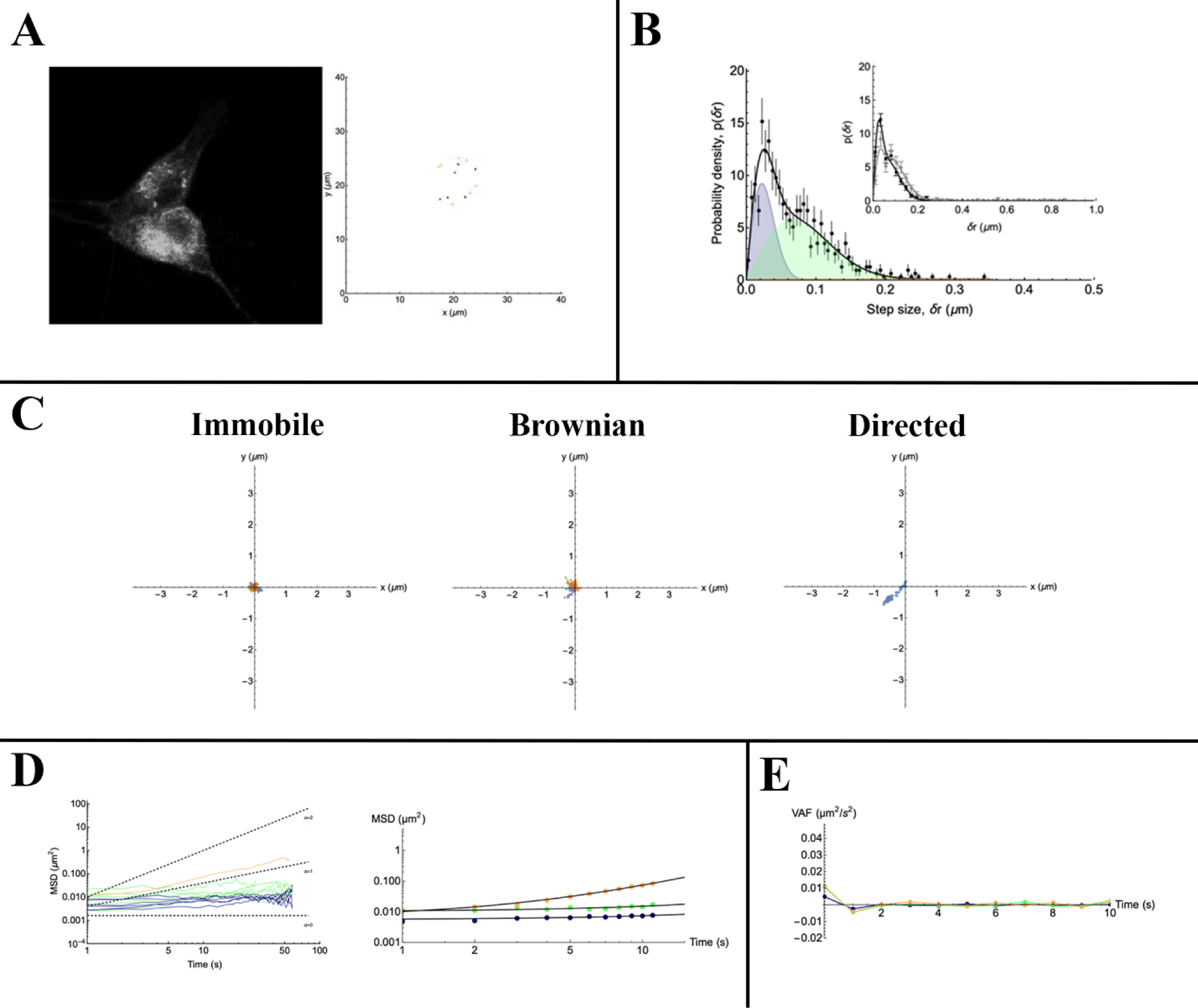
Trajectory analysis of EGFP-MamE particles. **A.** The confocal micrograph of a representative cell (left panel, black and white image of Figure 3A) is shown with trajectories of punctate particles detected within the cell (right panel). Immobile, Brownian, and directed trajectories are represented in black, green, and orange, respectively. **B.** The representative distribution of displacements is presented, where the immobile, Brownian, and direct populations are represented by the blue, green, and orange peaks, respectively. **C.** Immobile, Brownian, and directed trajectories are plotted on a cartesian plane. **D.** Immobile (black), Brownian (green), and direct (orange) trajectories are represented as mean-squared displacement (MSD) as a function of lag time. Dashed lines indicate the expected slope of the MSD, and the figure on the right shows average MSD for all trajectories. **E.** The plot shows average velocity autocorrelation function (VAF) of immobile (black), Brownian (green), and direct (orange) trajectories.

**Supplementary Figure S5:**
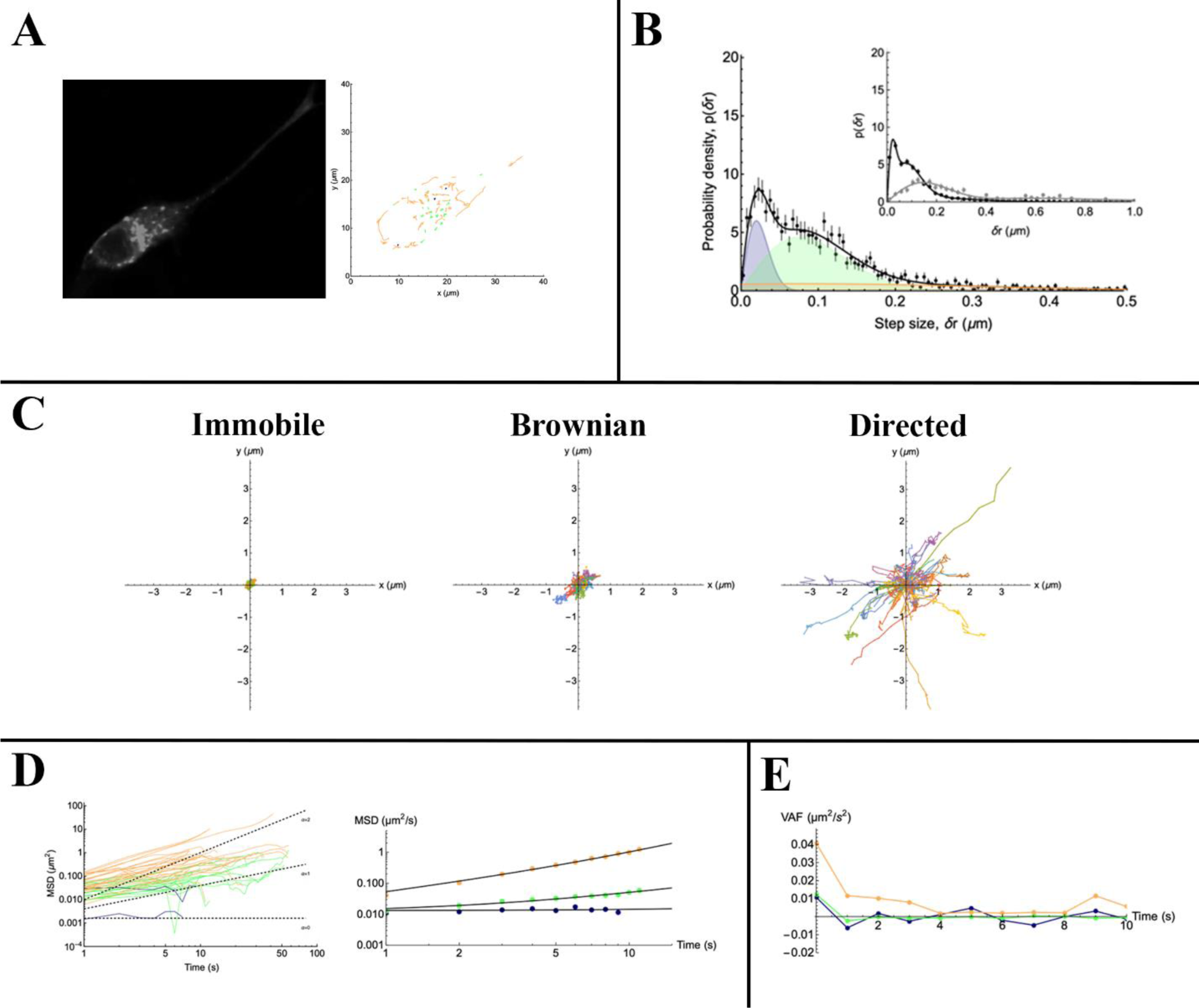
Trajectory analysis of Tomato-MamB particles. **A.** The confocal micrograph of a representative cell (left panel, black and white image of Figure 4A) is shown with trajectories of punctate particles detected within the cell (right panel). Immobile, Brownian, and directed trajectories are represented in black, green, and orange, respectively. **B.** The representative distribution of displacements is presented, where the immobile, Brownian, and direct populations are represented by the blue, green, and orange peaks, respectively. **C.** Immobile, Brownian, and directed trajectories are plotted on a cartesian plane. **D.** Immobile (black), Brownian (green), and direct (orange) trajectories are represented as mean-squared displacement (MSD) as a function of lag time. Dashed lines indicate the expected slope of the MSD, and the figure on the right shows average MSD for all trajectories. **E.** The plot shows average velocity autocorrelation function (VAF) of immobile (black), Brownian (green), and direct (orange) trajectories.

**Supplementary Figure S6:**
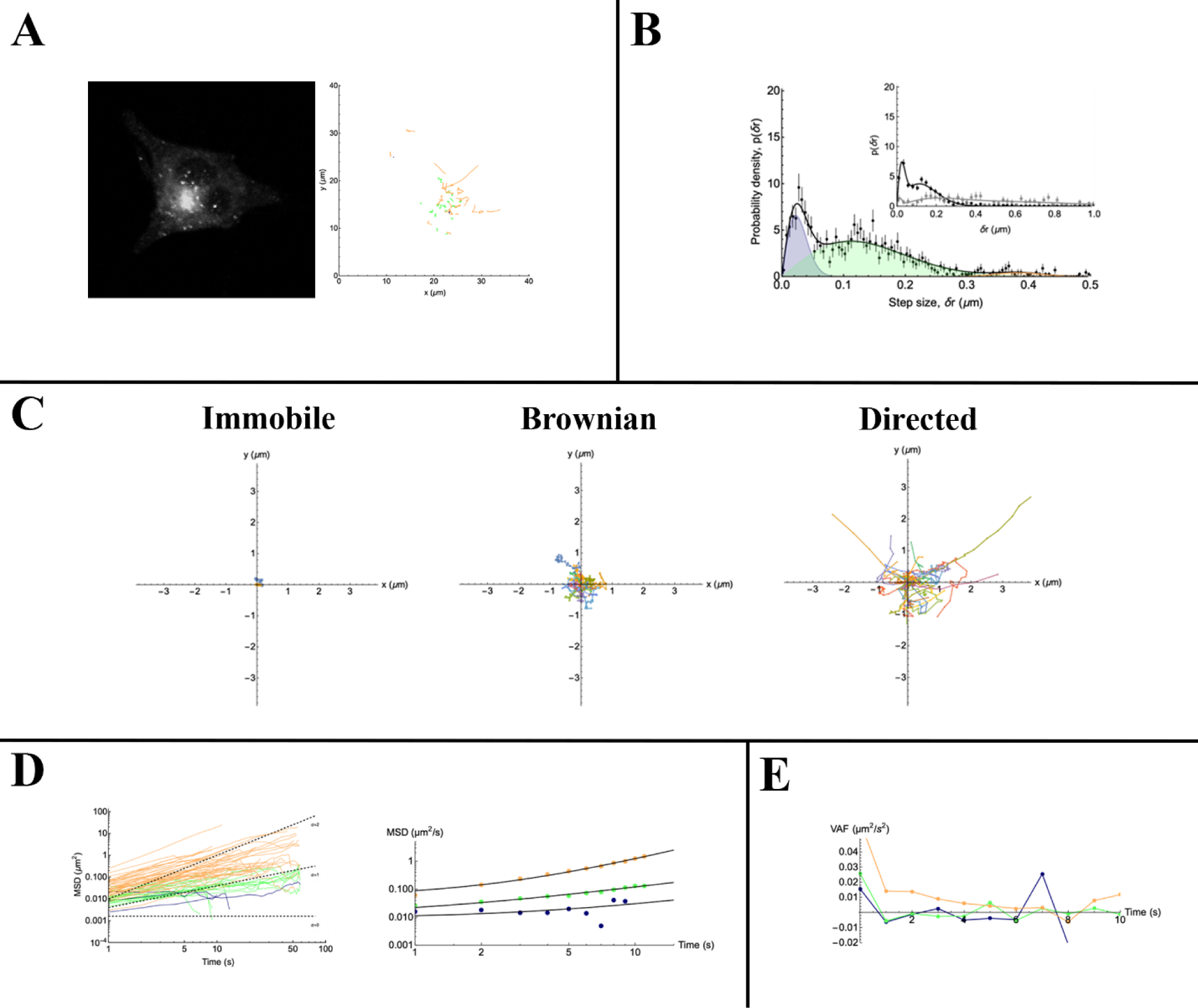
Trajectory analysis of Tomato-MamL particles. **A.** The confocal micrograph of a representative cell (left panel, black and white image of Figure 5A) is shown with trajectories of punctate particles detected within the cell (right panel). Immobile, Brownian, and directed trajectories are represented in black, green, and orange, respectively. **B.** The representative distribution of displacements is presented, where the immobile, Brownian, and direct populations are represented by the blue, green, and orange peaks, respectively. **C.** Immobile, Brownian, and directed trajectories are plotted on a cartesian plane. **D.** Immobile (black), Brownian (green), and direct (orange) trajectories are represented as mean-squared displacement (MSD) as a function of lag time. Dashed lines indicate the expected slope of the MSD, and the figure on the right shows average MSD for all trajectories. **E.** The plot shows average velocity autocorrelation function (VAF) of immobile (black), Brownian (green), and direct (orange) trajectories.

**Supplementary Figure S7:**
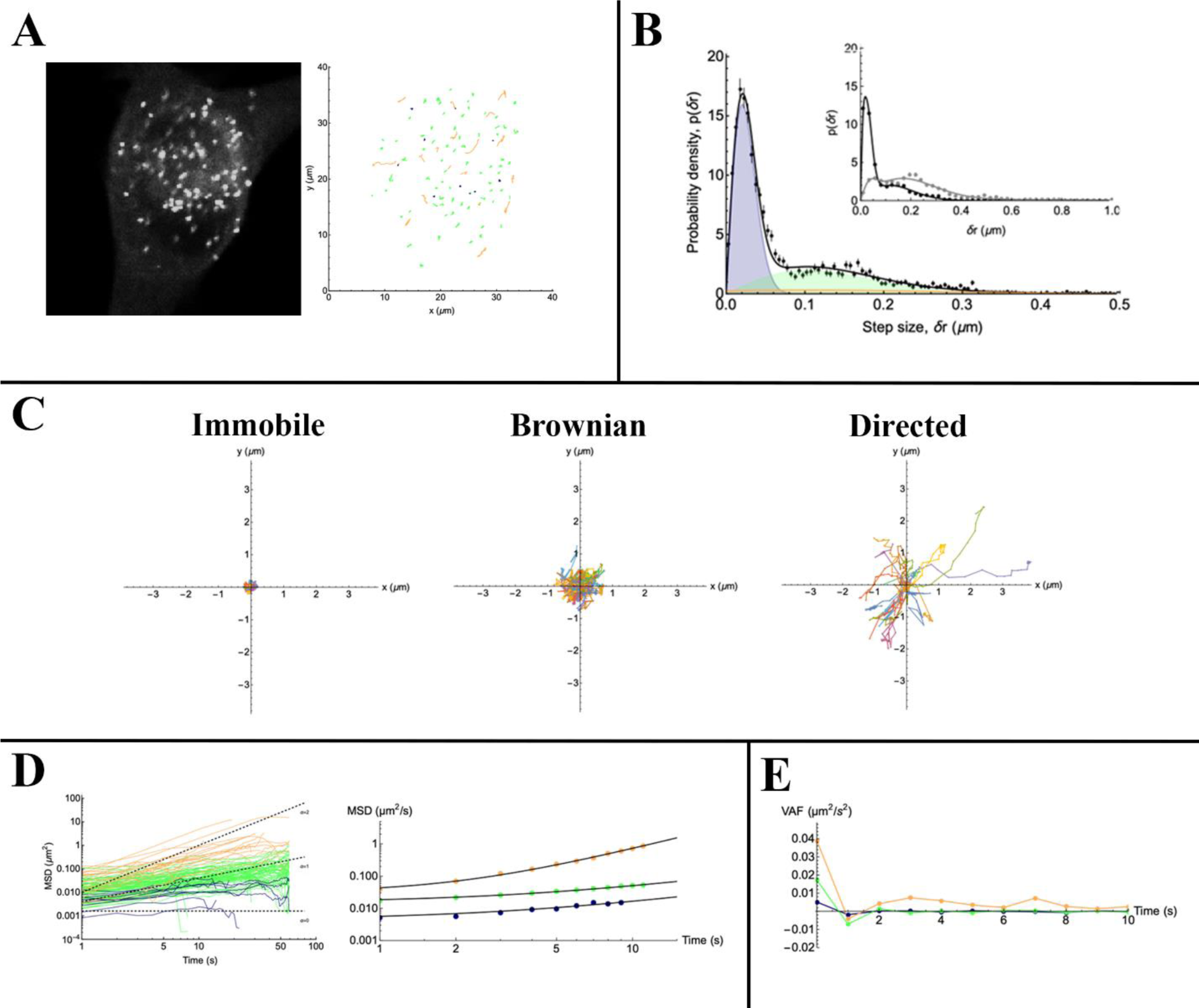
Trajectory analysis of Tomato-MamL_trunc_ particles. **A.** The confocal micrograph of a representative cell (left panel, black and white image of Figure 5D) is shown with trajectories of punctate particles detected within the cell (right panel). Immobile, Brownian, and directed trajectories are represented in black, green, and orange, respectively. **B.** The representative distribution of displacements is presented, where the immobile, Brownian, and direct populations are represented by the blue, green, and orange peaks, respectively. **C.** Immobile, Brownian, and directed trajectories are plotted on a cartesian plane. **D.** Immobile (black), Brownian (green), and direct (orange) trajectories are represented as mean-squared displacement (MSD) as a function of lag time. Dashed lines indicate the expected slope of the MSD, and the figure on the right shows average MSD for all trajectories. **E.** The plot shows average velocity autocorrelation function (VAF) of immobile (black), Brownian (green), and direct (orange) trajectories.

**Supplementary Figure S8:**
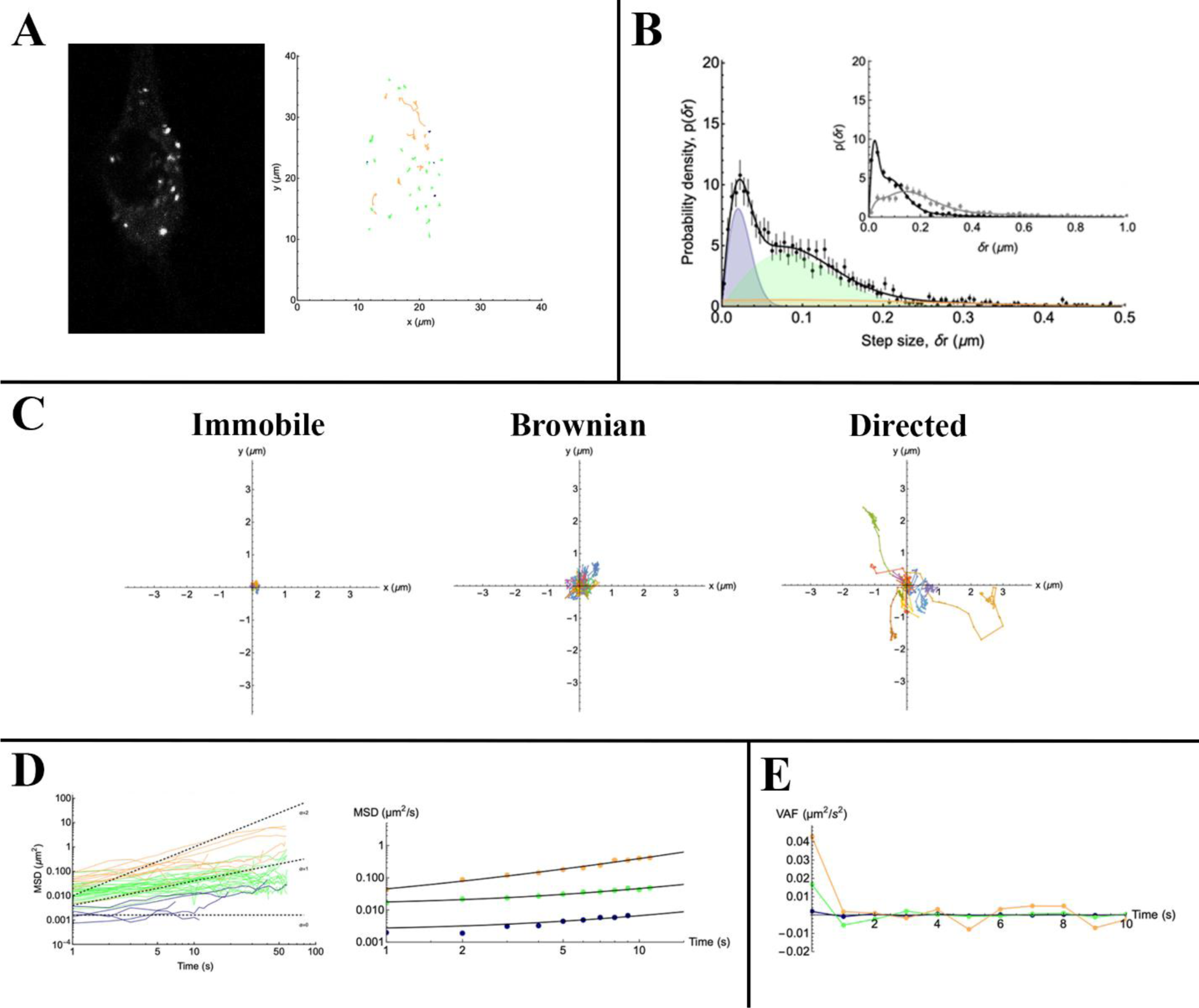
Trajectory analysis of Tomato-MamL/EGFP-MamI particles. **A.** The confocal micrograph of a representative cell (left panel, black and white image of Figure 6A) is shown with trajectories of punctate particles detected within the cell (right panel). Immobile, Brownian, and directed trajectories are represented in black, green, and orange, respectively. **B.** The representative distribution of displacements is presented, where the immobile, Brownian, and direct populations are represented by the blue, green, and orange peaks, respectively. **C.** Immobile, Brownian, and directed trajectories are plotted on a cartesian plane. **D.** Immobile (black), Brownian (green), and direct (orange) trajectories are represented as mean-squared displacement (MSD) as a function of lag time. Dashed lines indicate the expected slope of the MSD, and the figure on the right shows average MSD for all trajectories. **E.** The plot shows average velocity autocorrelation function (VAF) of immobile (black), Brownian (green), and direct (orange) trajectories.

**Supplementary Figure S9:**
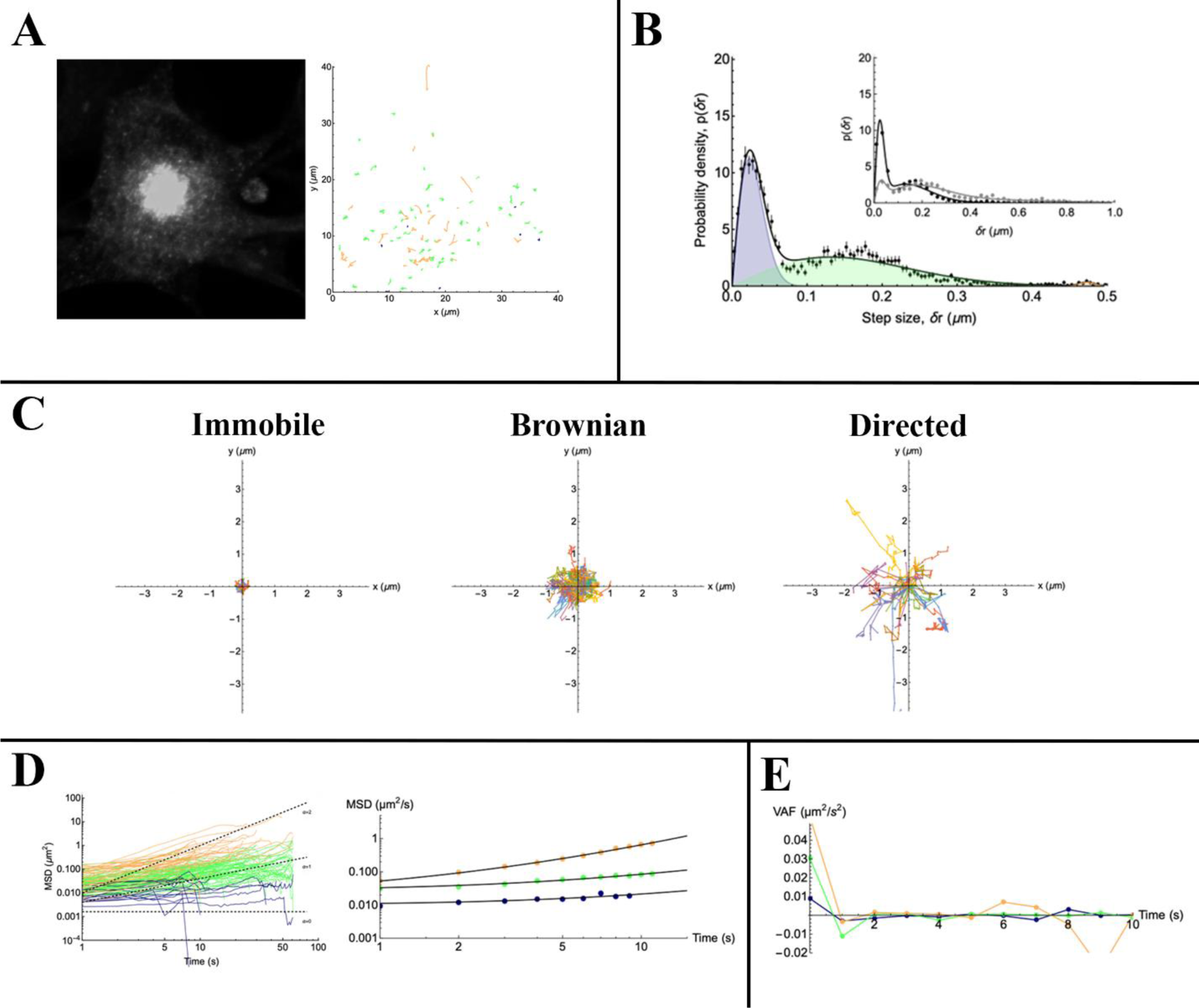
Trajectory analysis of FLAG-MamL/EGFP-MamI particles. **A.** The confocal micrograph of a representative cell (left panel, black and white image of Figure 6D) is shown with trajectories of punctate particles detected within the cell (right panel). Immobile, Brownian, and directed trajectories are represented in black, green, and orange, respectively. **B.** The representative distribution of displacements is presented, where the immobile, Brownian, and direct populations are represented by the blue, green, and orange peaks, respectively. **C.** Immobile, Brownian, and directed trajectories are plotted on a cartesian plane. **D.** Immobile (black), Brownian (green), and direct (orange) trajectories are represented as mean-squared displacement (MSD) as a function of lag time. Dashed lines indicate the expected slope of the MSD, and the figure on the right shows average MSD for all trajectories. **E.** The plot shows average velocity autocorrelation function (VAF) of immobile (black), Brownian (green), and direct (orange) trajectories.

**Supplementary Figure S10:**
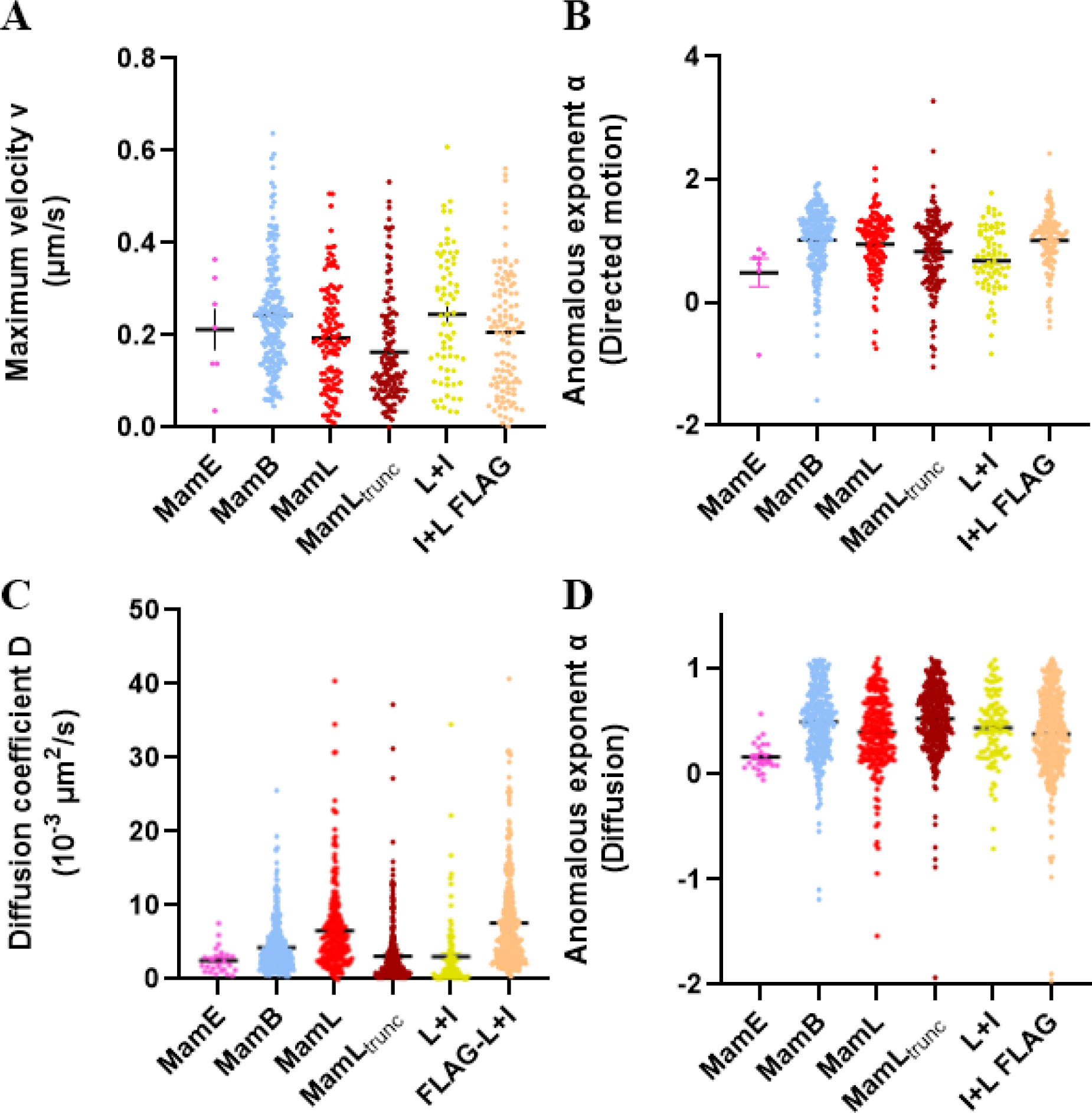
Comparison of trajectory parameters for individual particles detected in cells expressing magnetosome proteins. In trajectories classified as directed motion, scatter plots indicate the range in both maximum velocity (A) and anomalous exponent (B). Likewise, in trajectories classified as entirely Brownian, the range in both diffusion coefficients (C) and anomalous exponents (D) were extracted.

**Table S1.**
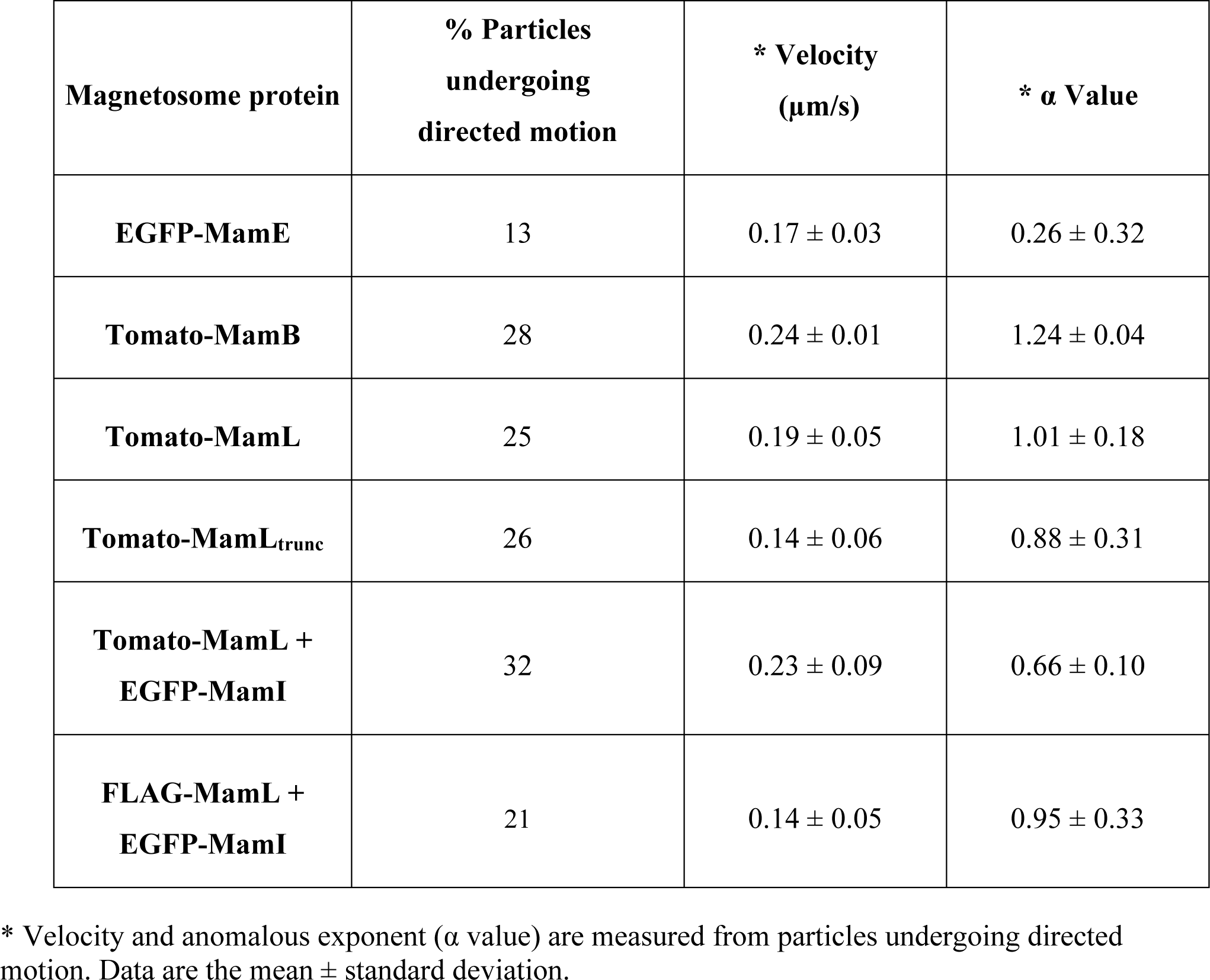
Summary of velocity and anomalous exponent values for particles undergoing directed motion.

| **Magnetosome protein** | **% Particles undergoing directed motion** | **\* Velocity ($\mu\text{m/s}$)** | **\* $\alpha$ Value** |
| --- | --- | --- | --- |
| **EGFP-MamE** | 13 | $0.17 \pm 0.03$ | $0.26 \pm 0.32$ |
| **Tomato-MamB** | 28 | $0.24 \pm 0.01$ | $1.24 \pm 0.04$ |
| **Tomato-MamL** | 25 | $0.19 \pm 0.05$ | $1.01 \pm 0.18$ |
| **Tomato-MamL<sub>trunc</sub>** | 26 | $0.14 \pm 0.06$ | $0.88 \pm 0.31$ |
| **Tomato-MamL + EGFP-MamI** | 32 | $0.23 \pm 0.09$ | $0.66 \pm 0.10$ |
| **FLAG-MamL + EGFP-MamI** | 21 | $0.14 \pm 0.05$ | $0.95 \pm 0.33$ |

\* Velocity and anomalous exponent ( $\alpha$ value) are measured from particles undergoing directed motion. Data are the mean ± standard deviation.

